# Hierarchical organization of developing HSPC in the human embryonic liver

**DOI:** 10.1101/321877

**Authors:** Y. Zhang, D. Clay, M.T. Mitjavila-Garcia, A. Alama, B. Mennesson, H. Berseneff, F. Louache, A. Bennaceur-Griscelli, E. Oberlin

**Affiliations:** Inserm, UMR 1170, Villejuif, France; Paris-Saclay University, Villejuif, France; Gustave Roussy, Villejuif, France; Inserm UMS 33, Villejuif, France; Inserm UMR 935, Villejuif, France; André Lwoff Institute (IFR89), Villejuif, France; René-Dubos Hospital, Obstetrics and Gynecology Department, Pontoise, France

**Keywords:** human embryo, hematopoietic stem and progenitor cell, embryonic liver, VE-cadherin, Angiotensin-converting enzyme

## Abstract

Despite advances to engineer transplantable hematopoietic stem and progenitor cells (HSPCs) for research and therapy, an in depth characterization of the developing human hematopoietic system is still lacking. The human embryonic liver is at the crossroad of several hematopoietic sites and harbours a complex hematopoietic hierarchy including the first, actively dividing, HSPCs that will further seed the definitive hematopoietic organs. However few is known about the hierarchical phenotypic and functional hematopoietic organization operating at these stages of development.

Here, by using a combination of four endothelial and hematopoietic surface markers i.e. the endothelial-specific marker VE-cadherin, the pan-leukocyte antigen CD45, the hemato-endothelial marker CD34 and the Angiotensin-Converting Enzyme (ACE, CD143), encompassing all early human HSPCs, we identified a hematopoietic hierarchy and, among it, a population co-expressing the four markers that uniquely harbored a proliferation and differentiation potential both *ex vivo* and *in vivo*. Moreover, we traced back this population to the yolk sac and AGM sites of hematopoietic emergence. Taken together, our data will help to identify human HSPC self-renewal and amplification mechanisms for future cell therapies.

**SUMMARY STATEMENT:** We uncover the phenotypic and functional hematopoietic hierarchy operating in the early human embryo. It will bring insights into the mechanisms driving hematopoietic stem cell self-renewal for future cell therapies.

## INTRODUCTION

Hematopoietic stem and progenitor cell (HSPC) transplantation is an important therapy used for over 40 years for patients suffering hematological malignancies. However the demand for clinical grade allogeneic HSPCs has significantly increased over the past decades with no parallel increase of the HSPC offer. A major goal in regenerative medicine is to derive HSPCs from several nonhematopoietic sources and manipulate them to treat patients. Despite many efforts, the generation of transplantable HSPCs from pluripotent stem cells or from directly reprogrammed cells is not yet possible. A similar remark applies for the conditions of HSPC amplification *ex vivo*.

These questions are calling for additional studies to identify the founders of the human hematopoietic system and the mechanism underlying HSPC emergence and amplification.

During embryonic development, the first hematopoietic cells (HCs) emerge in the yolk sac (YS) and generate primitive erythroblasts, macrophages and megakaryocytes (Tober et al., 2007; Palis et al., 1999; Silver and Palis, 1997; Luckett et al., 1978). This first wave is followed by a second wave of erythro-myeloid progenitors also derived from the YS that gives rise to definitive erythroid, megakaryocyte, myeloid, and multipotent progenitors (Palis and Yoder, 2001; Palis et al., 1999, Migliaccio et al., 1986) that seed the fetal liver and contribute to the generation of erythropoietic progenitors prior to HSPC colonization.

The first HSPCs emerge from specialized endothelial cells (EC), known as hemogenic endothelium (HE), in the aorta-gonad-mesonephros (AGM) region, migrate to the fetal liver to be massively amplified and finally reach the bone marrow (BM) their life-long residence (Ivanovs et al., 2017; Klaus and Robin, 2017; Crisan and Dzierzak, 2016; Ciau-Uitz and Patient, 2016; Julien et al., 2016; Medvinsky et al., 2011). This conversion of endothelium into HCs termed endothelial-to-hematopoietic transition (EHT) has been extensively verified in various animal model (Boisset et al., 2015 and 2010; Chen et al., 2011 and 2009; Bertrand et al., 2010; Kissa and Herbomel, 2010; Eilken et al., 2009; Zovein et al., 2008; Fraser et al., 2002; Jaffredo et al., 1998; Nishikawa et al., 1998a) and we were among the first to probe this blood-forming potential in the human YS, AGM and Embryonic Liver (EL) (Oberlin et al., 2002). These AGM-derived HSPCs are the founders of the definitive hematopoietic system. They display a dual endo-hematopoietic phenotype characterized by the expression of the endothelial-specific junctional vascular endothelial cadherin (Cdh5, CD144, hereafter designated as VEC) and the presence of the pan-leukocyte antigen CD45 (Ivanovs et al., 2014; Rybtsov et al., 2014 and 2011; Taoudi et al., 2008; Fraser et al., 2003; North et al., 2002; Nishikawa et al., 1998b).

Since HSPCs produced in the AGM are likely to seed the EL, we hypothesized several years ago that cells expressing both endothelial and hematopoietic traits could also be detected in the human EL, in keeping with the situation described in the mouse fetal liver (Kim et al., 2005, Taoudi et al., 2005). We indeed identified a sub-population of HSPCs in the human EL that co-expressed the endothelial-specific marker VEC, the pan-leukocyte antigen CD45 and the hemato-endothelial marker CD34 and we named this population 34DP since it expressed the CD34 antigen and was double positive (DP) for VEC and CD45. We showed that the 34DP cell fraction represented an intermediate between the hemogenic ECs that we previously identified in the human embryo (Oberlin et al., 2002) and the more advanced HC population that lacked VEC expression present in the late embryonic and fetal liver hematopoietic stages (Oberlin et al., 2010a and 2010b). This 34DP subset displayed important self-renewal and differentiation capacities, as detected by *ex vivo* and *in vivo* hematopoietic assays compared to its VEC negative counterpart. The 34DP population thus represents a potential model to study the earliest HSPCs in the human embryo and the mechanisms operating during their amplification and maturation. However the precise hematopoietic hierarchy operating at these stages was not known.

We report here an ensemble of phenotypic and functional characterizations (Figure 1) that allows hierarchically define the subpopulations present in the human EL hematopoietic HSPC compartment. We used markers that typifies the most primitive HSPCs during adult (Majeti et al., 2007) and embryonic stages (Sinka et al., 2012; Ivanovs et al., 2017), and in particular, the CD143 transmembrane molecule, also designated as Angiotensin-Converting Enzyme (ACE), identified as a cell surface marker of adult HSPCs (Jokubaitis et al., 2008) and known to be expressed on the earliest HSPCs in developing blood-forming tissues of the human embryo and fetus (Sinka et al., 2012). We demonstrated that VEC, CD45 and CD34 in conjunction with ACE allows to separate several subpopulations endowed with contrasted hematopoietic potential and to identify a subpopulation uniquely able to maintain a proliferative potential and an undifferentiated state *ex vivo* and *in vivo*. Moreover, we also tested whether other hematopoietic territories earlier in the developing human conceptus harbored cells with the 34DP signature and confirm that these cells can be identified in sites of HSPC emergence, including the YS, the AGM region and to a much lesser extend the placenta. In contrast, neither the cord blood nor the adult bone marrow carried cells with the 34DP signature. This work will help to identify the molecular mechanisms involved in the first human HSPC self-renewal and amplification to improve future therapies.

**Figure 1.**
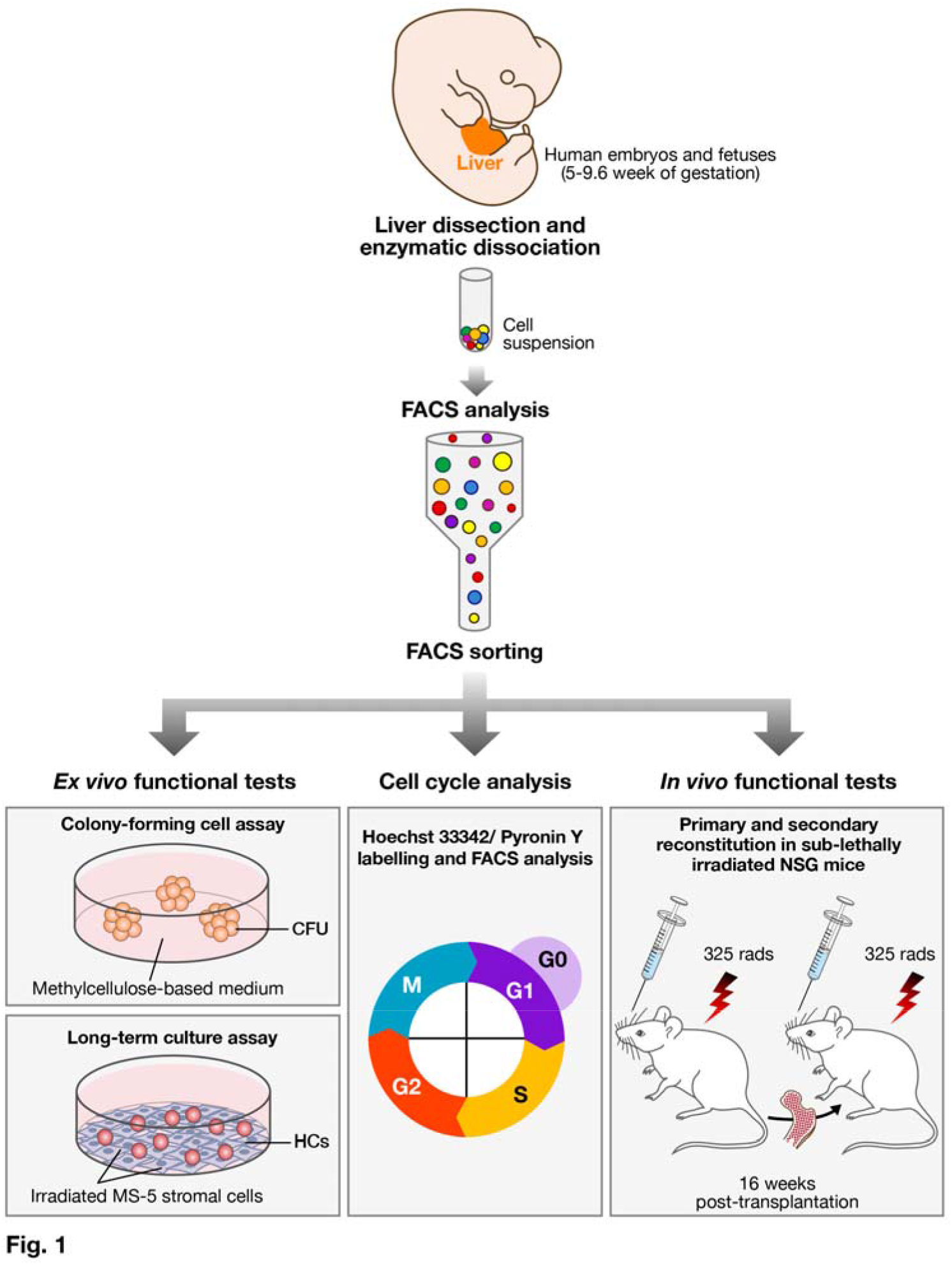
Experimental protocol for embryonic and fetal liver cell populations analysis.

Embryonic and fetal livers were excised and enzymatically and mechanically dissociated. Recovered cells were FACS analyzed and selected populations were cell sorted. Sorted cells were analysed either *ex vivo* by performing colony-forming cell-assays, long-term culture assays and cell cycle experiments or *in vivo* by challenging the primary and secondary reconstitution potential in NSG mice.

## MATERIALS AND METHODS

### Human tissues

Human embryos and fetuses from 5 to 9.6 weeks of gestation were obtained following voluntary abortions. Tissue collection and use were performed according to the guidelines and with the approval of the French National Ethic Committee (authorization N° PFS10_011). Developmental age was estimated based on several anatomic criteria according to the Carnegie classification for embryonic stages (O’Rahilly et al., 1987) and by ultrasonic measurements for fetal stages. Supplemental Table S1 recapitulates the stages of the embryonic and fetal tissues used in this study, as well as the type of experiments performed for each type of tissue.

Human umbilical cord blood (CB) samples were collected in citrate phosphate dextrose solution from healthy full-term newborns. CB samples were obtained through a partnership with the Cord Blood Bank of St Louis Hospital (Paris, France) which is authorized by the French Regulatory Authority (authorization N° PPC51).

Human bone marrow (BM) samples were collected from healthy individuals undergoing hip surgery (CHR Robert Ballanger hospitals, France).

All human samples were obtained with the written informed consents of subjects according to Helsinki declaration.

### Cell preparation

Embryonic and fetal organs were excised sterilely using microsurgery instruments and a dissecting microscope, in phosphate-buffered saline (PBS) containing 100 U/ml penicillin and 100 mg/ml streptomycin (Invitrogen, Cergy-Pontoise, France). Tissues were dissociated for 1 hour at 37°C in Iscove’s modified Dulbecco medium (IMDM; Invitrogen) containing 10% fetal calf serum (FCS; Hyclone Laboratories, Logan, UT), 0.1% type I/II/IV collagenase and type VIII hyaluronidase (Sigma-Aldrich, St Quentin Fallavier, France). Tissues were then disrupted mechanically through 18-, 23- and 26-gauge needles successively. Cell clumps were removed on a 50-μm nylon mesh (BD Biosciences, Le Pont de Claix, France).

CB and BM mononuclear cells (MNCs) were isolated by a Pancoll (Dutscher, Brumath, France) density gradient centrifugation.

All recovered cells were counted, after red blood cell lysis in 0.1% acetic acid. Cell viability was determined by 0.2% trypan blue exclusion (Invitrogen).

### Sorting and cell analysis by flow cytometry

Monoclonal antibodies (Mabs) used for cell sorting or FACS analysis are listed in Supplemental Table S2. For cell sorting, cells were incubated for 30 min on ice with FITC-anti-CD45, PE-anti-VEcadherin, APC-anti-CD143 and APC-Alexa Fluor 750-anti-CD34 antibodies. Labeled cells were washed in PBS (Invitrogen) 0.2% bovine serum albumin (BSA) (Sigma-Aldrich), re-suspended in complete IMDM medium (Invitrogen), 10% fetal calf serum (FCS; Hyclone Laboratories), 100 U/ml penicillin, 100 mg/ml streptomycin and 2 mM L-glutamine (Invitrogen). Selected populations were sorted on a FacsDiva cell sorter (BD Biosciences). Sorted cells were systematically reanalyzed to establish purity.

Cells developed in culture were harvested by non-enzymatic treatment (Cell dissociation solution; Sigma-Aldrich), washed in complete medium, labeled with fluorochrome-conjugated antibodies to hematopoietic and endothelial markers and analyzed on a FACSCalibur flow cytometer (BD Biosciences) using the CellQuest^TM^ software (BD Biosciences).

BM cells collected from primary and secondary transplanted NSG mice were washed, labeled with fluorochrome-conjugated antibodies specific to human hematopoietic differentiation. Staining with lineage specific antibodies was performed concurrently with anti-human CD45 and anti-murine CD45 Mabs. Specificity of human Mabs was checked on BM cells from non-transplanted mice. Stained cells were analyzed on a FACScanto II cell analyser (BD Biosciences) using the FACSDiva^TM^software (BD Biosciences).

In all cases, background staining was evaluated using isotype-matched control antibodies and 7-Amino-Actinomycine D (7AAD) (Sigma-Aldrich) was used to gate dead cells out.

### Cell culture

#### - Colony-forming cell (CFC) assays

FACS sorted cells were plated in methylcellulose-based medium (Methocult H4434, StemCell Technologies) according to the manufacturer instructions. After 12 days of culture, colonies (≥ 50 cells) were scored and photographed using an Olympus IM inverted microscope (Olympus) equipped with a Coolpix 5400 camera (Nikon). Large colonies > 0.5 mm in diameter were scored as high proliferative-potential colony-forming unit (HPP-CFU) colonies, while smaller colonies <0.5 mm were considered as low proliferative-potential colony-forming unit (LPP-CFU) colonies. After counting, colonies were harvested from methylcellulose, dispersed to single cell suspension and washed in complete IMDM medium. Recovered cells were labeled with fluorochrome-conjugated antibodies to hematopoietic differentiation for FACS analysis and cell counting. Cell number fold increase between day 0 and day 12 of culture was obtained by calculating the ratio between the quantity of cells obtained at day 12 and the quantity of cells seeded at day 0.

#### - Long-term culture-initiating cell assay on MS-5 stromal cells

Sorted cells were cultivated in bulk on MS-5 mouse bone marrow stromal cells (Itoh et al., 1989) as previously described (Tavian et al., 2001), except that only 3 human recombinant cytokines were added: 50 ng/ml stem cell factor, 1 ng/ml interleukin-15 and 5 ng/ml interleukin-2 (AbCys, Paris, France). Half of the medium was replaced weekly. Every week, cells collected from wells with significant proliferation were washed, counted in 0.2% trypan blue and FACS analyzed after labeling with fluorochrome-conjugated antibodies to hematopoietic differentiation. Cell number fold increase was obtained by calculating the ratio between the number of cells obtained at day 25 and 35 of the co-culture and the quantity of cells seeded at day 0. Colonies of HCs were scored and photographed under an ICM-405 phase-contrast inverted microscope (Carl Zeiss) using an XC-003 3CCD color camera (Sony) at different time point of the co-culture.

### Analysis of HC division by 5,6-Carboxyfluorescein-Diacetate-Succinimidyl-Ester (CFSE) dye dilution

HCs labeling by CFSE was carried out as described previously (Glimm and Eaves, 1999). Briefly, cells were grown on MS-5 stromal cells and incubated with 2.5 mM CFSE (Molecular Probes^TM^, Thermo Fischer Scientific, Villebon sur Yvette, France) for 10 minutes at 37°C and then quenched with cold FCS. After 3 washes in 10% FCS complete medium, labeled cells were then co-cultured with MS-5 cells for 3 and 6 additional days. The percentage of proliferating cells was analyzed by flow cytometry measurements of the dilution of CFSE on day 3 and 6 after staining with an anti-CD45-APC mAb using a FACSCalibur flow cytometer (BD Biosciences) and the FlowJo^TM^ software’s proliferation tool (BD Biosciences). Cells undergoing division were identified by the decrease in CFSE resulting from dilution of the dye with each division. The number of cell divisions was calculated between day 0 undivided cells (CSFE bright) and the first peak obtained at day 3 and day 6.

### Cell cycle analysis

Quiescent (G_0_) and cycling (G_1_ and S + G_2_/M) cells were separated based on Hoechst/Pyronin-Y fluorescence as described (Shapiro, 1981). Briefly, 10.10^6^ EL cells per ml were suspended in prewarmed Dulbecco’s modified Eagle’s medium, 2% FCS, 10 mM HEPES, Hoechst 33342 (final concentration: 10 mg for 10.10^6^ cells per ml) (all from Invitrogen) and were incubated at 37°C for 30 min. Pyronin-Y (Sigma-Aldrich; final concentration: 300 ng for 10.10^6^ cells per ml) was added for 15 min at 37°C at the end of Hoechst incubation. After incubation, cells wer**e** washed and suspended in 0.2% BSA cold PBS, immunostained with FITC-anti-VE-cadherin, BV421-anti-CD45, APC-anti-CD143 and APC-Alexa Fluor 750-anti-CD34 Mabs. Labeled cells were washed in 0.2% BSA cold PBS and analyzed using a LSR Fortessa^TM^ flow cytometer (BD Biosciences) and the FlowJo^TM^ software’s cell cycle tool.

### Hematopoietic repopulating activity in NSG mice

NOD/shi-SCID IL2Rγ^-/-^ (NSG) mice were purchased from the Jackson laboratory and were bred and maintained under specific pathogen free conditions at the animal facility of Gustave Roussy Institute (Permit n° E-94-076-11). Animal experiments were conducted in accordance with institutional and national guidelines and were approved by Ethical Committee C2EA-26 officially registered with the french ministry of research (Permit n°: 2012-148).

Male or female mice, 6 to 8 week-old, were irradiated at 3.5 Gy with an IBL637 gamma irradiator containing a Cs-137 source (CIS Bio International, Saclay, France). Mice were intravenously injected 8 to 24 hours after irradiation with sorted cells. Human HC engraftment was assessed 16 weeks post-transplantation by multicolor flow cytometry analysis as described below. For secondary transplantation, the cellular content of two femur and two tibias was flushed and human cells were isolated by EasySep^TM^Mouse/Human chimera isolation kit (Stemcell Technologies, Grenoble, France). Isolated human cells were intravenously injected into irradiated mice. Human HC engraftment was assessed 16 weeks later by multicolor flow cytometry analysis as described below.

### Statistical Analysis

Data are representative of at least three independent experiments performed on different embryos and are expressed as mean ± standard deviation (SD). Statistical significance was determined using the non-parametric Mann-Whitney U test. *P*<0.001^***^ was considered highly significant.

## RESULTS

### Phenotypic characterization of VE-cadherin^+^ CD45^+^ double positive (DP) population in the human EL

To document the hematopoietic hierarchies operating in the early human embryo, we searched for markers that would accurately identify recently emerged HSPCs in the EL. We first addressed the value of several markers used to identify human HSPC (Majeti et al., 2007) such as CD90 and CD117 (also known as c-KIT), CD38 and CD45RA. We also tested CD105 (also known as Endoglin), a well-known endothelial marker (Cheifetz et al., 1992). All markers were used in combination with VEC, CD45 and CD34 antigens to estimate their added value in the DP (VECAD^low^CD45^low^) cells and to compare DP to EC (VECAD^+^ CD45^-^) and HC (VECAD^-^CD45^high^ and VECAD^-^CD45^low^) EL cell populations. ELs were microdissected from embryos ranging from 5 to 9.6 weeks and dissociated by collagenase digestion and mechanical disruption and further processed for flow cytometry analysis and cell sorting (Figure 1 and Supplemental Table S1).

Flow cytometry analysis confirmed the existence of VEC^+^ CD45^-^ ECs, VEC^low^CD45^low^ DP cells and VEC^-^CD45^low^ and VEC^-^CD45^high^ HC populations (Figure 2A). These populations were found along the different EL developmental stages examined albeit at different frequencies i.e., DP cells decreased with time ranging from 3.92% to 0.18% between 5 and 9.6-weeks of gestation (Supplemental Figure S1). VEC expression was also found in more differentiated CD45^high^ cells (VEC^+^ CD45^high^) (Figure 2A), likely monocytes, that expressed a low level of CD34 and myeloid lineage markers (data not shown). A large number of DP cells were positive for the HSPC markers CD34, CD90 and CD117 and were negative for CD38 and CD45RA, a phenotype likely associated with primitive progenitors, including stem cells. Conversely, most of the cells in the HC fractions were negative for the HSPC markers and positive for CD38 and CD45RA. CD105, a typical endothelial marker at that stage, was also found in DP but not in HCs hence confirming the endothelial signature of the DP population (Supplemental Table S3).

**Figure 2.**
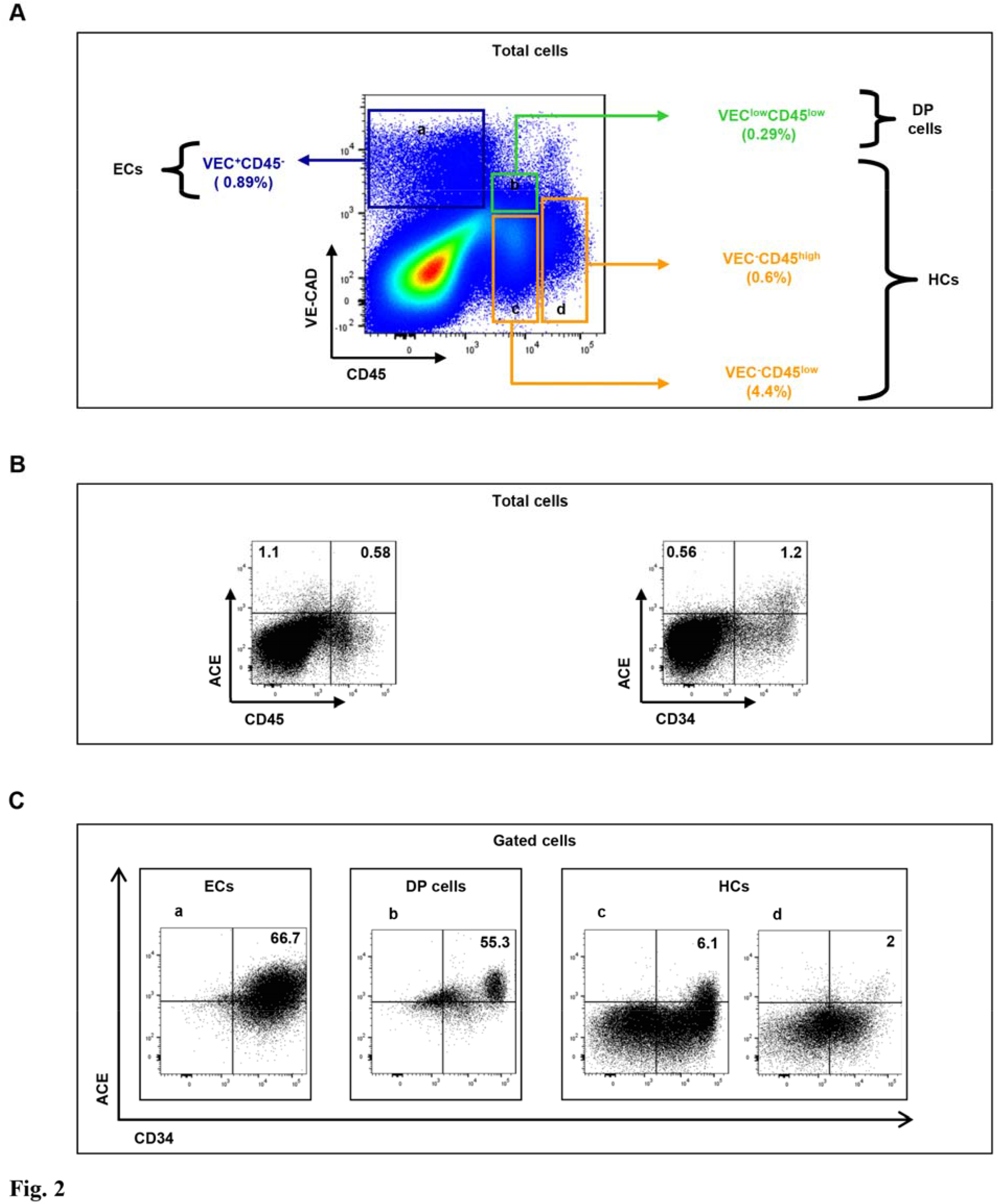
CD45, CD34, VE-cadherin and ACE FACS analysis on human EL cells.

Whole liver cell from a 7.5-week-old human embryo quadruple-stained with anti-CD45, -VEC, -CD34 and -ACE Mabs.

**(A)** Flow cytometry analysis delineating VECAD^+^ CD45^-^ endothelial (frame **a)**, VECAD^low^CD45^low^ DP (frame **b**) and VECAD^-^CD45^high^ (frame **c**), VECAD^-^CD45^low^ hematopoietic (frame **d**) EL cell populations.

**(B)** Flow cytometry analysis showing relative expression of CD45, CD34 and ACE in total EL cells. **(C)** Flow cytometry analysis showing relative expression of CD34 and ACE by VEC^+^ CD45^-^ endothelial (frame **a**), VEC^low^CD45^low^ DP (frame **b**) and VEC^-^CD45^low^ (frame **c**), VEC^-^CD45^high^ (frame **d**) hematopoietic EL cell populations **(A, B, C)**: Numbers indicate the percentages of positive cells in the corresponding frames.

We also addressed the value of the angiotensin-converting enzyme (ACE, CD143) in combination with VEC, CD45 and CD34 antigens in the ECs, HCs and DP cells. Whole liver cell analysis demonstrated that ACE was expressed in both CD45^+^ and CD45^-^ cell population whereas it was mostly expressed in CD34^+^ cells (Figure 2B). A large number of VEC^+^ CD45^-^ ECs (66.7±11%; Figure 2Ca) and VEC^low^CD45^low^ DP cells (70.2±9%; Figure 2Cb) expressed ACE while the HC fraction also contained ACE-expressing cells, but with a lower frequency (6.1±4% and 2±0.78% in VECAD^-^ CD45^low^ and VECAD^-^CD45^high^ cells respectively; Figure 2Cc-d) and mean fluorescence intensity (MFI) compared to EC and DP cells. Because ACE was the marker the most tightly associated with DP cells, we decided to include it to our VEC, CD45 and CD34 antigen panel to characterize more deeply the EL HSCP hierarchy.

### *Ex vivo* expansion ability is restricted to the VEC^low^CD45^low^CD34^high^ACE^+^ EL cell subset

To test the self-renewal and multilineage hematopoietic potential in the EL hematopoietic populations, we performed colony-forming cell (CFC) and long-term culture assays (Figure 1). In both assays the hematopoietic potential of VEC^low^CD45^low^CD34^high^ACE^+^ (34DPACE^+^) cell population was compared to that of VEC^low^CD45^low^CD34^low^ACE^-^ (34LDPACE^-^), VEC^-^CD45^low^CD34^high^ACE^+^ (34HCACE^+^), VEC^-^CD45^low^CD34^high^ACE^-^ (34HCACE^-^) and VEC^-^CD45^low^CD34^low^ACE^-^(34LHCACE^-^) cell populations sorted from the same EL according to the sorting strategy depicted in Supplemental Figure S2.

#### CFC-assays

The presence of hematopoietic progenitors in each sorted cell population was first estimated by CFC assay. All cell populations contained hematopoietic progenitors, except 34LDPACE^-^ cells. 34LHCACE^-^ produced very few, small, colonies and we did not further consider it. The greatest number of colonies was found in the 34HCACE ^+^ and 34HCACE^-^ cell populations compared to the 34DPACE ^+^ cell population (233±10, 176.5±21 and 130±14 CFCs respectively) (Figure 3A). However, the 34DPACE ^+^ cell population contained exclusively CFCs with high proliferative potential, whereas the 34HCACE^-^ population only contained CFCs with low proliferative potential. 34HCACE ^+^ cells stood between the 34DPACE ^+^ and 34HCACE^-^ cell populations and contained more CFC with low proliferative potential than high proliferative potential (131.5±26 and 45±6 respectively) (Figure 3A).

**Figure 3.**
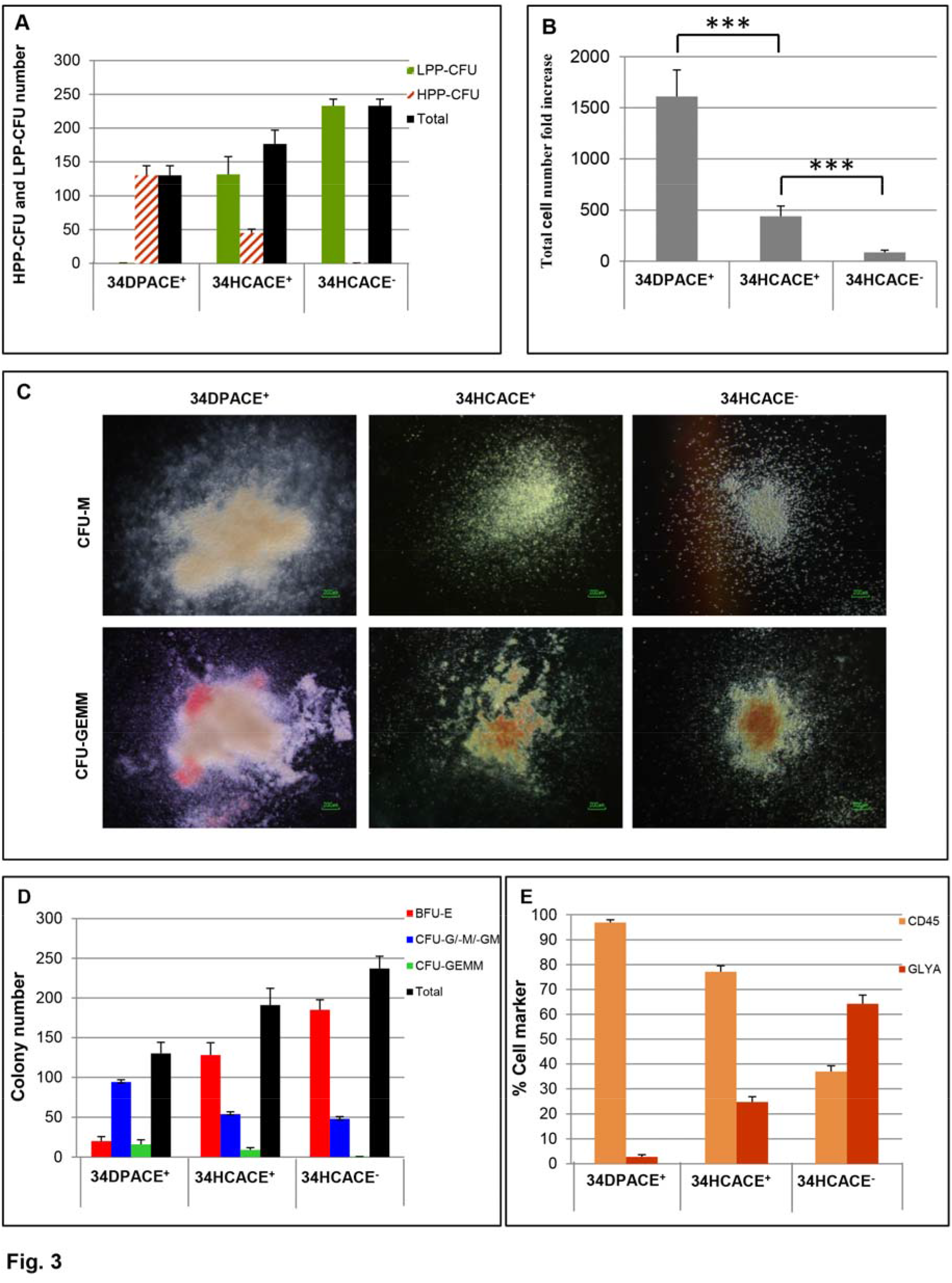
Clonogenic potential of 34DPACE ^+^,34HCACE^+^ and 34HCACE^-^ EL cell fractions. **(A)** High- (HPP) and low- (LPP) CFC numbers normalized to 1.10^4^ cells for each sorted EL cell population. **(B)** Total cell number fold increase for each sorted EL cell populations. **(D)** BFU-E, CFU-G/M/GM and CFU-GEMM colony numbers normalized to 1.10^4^ cells for each sorted EL cell population. **(E)** Flow cytometry analysis after CD45 and Glycophorin A staining for each sorted EL cell populations. (**A, B, D, E**): results show mean values of triplicates with standard deviation in one representative experiment from three independent experiments performed on 6.8 to 8.1-week-old ELs. HPP-CFU, high proliferative-potential colony-forming unit; LPP-CFU, low proliferative-potential colony-forming unit; BFU-E, Burst forming unit-erythroid; CFU-G/M/GM, colony forming unit-granulocyte, -macrophage, -granulocyte-monocyte; CFU-GEMM, colony forming unit -granulocyteerythrocyte-macrophage-megakaryocyte; GLYA, Glycophorin A. p-value summary (Mann-Whitney U test): ^***^p < 0.001.

The proliferative capacity of CFCs cells was evaluated in each cell population by measuring the cell number fold-increase between day 0 and 12 of culture. The amplification of 34DPACE ^+^ cells was 4 fold higher than that of 34HCACE ^+^ and 20 fold higher than that of 34HCACE^-^ cells (Figure 3B) hence revealing a strong proliferative capacity. Figure 3C shows typical CFU-GM and CFU-GEMM colonies obtained from each sorted cell populations and confirmed that colonies produce from 34DPACE ^+^ cells were always larger and more densely populated than colonies obtained from 34HCACE ^+^ or 34HCACE^-^ cell populations.

Hematopoietic lineage distribution within colonies showed that 34DPACE ^+^ cell produced more CFUG/M/GM colonies than BFU-E colonies (94±3 vs 20±6 respectively) whereas 34HCACE ^+^ and 34HCACE^-^ cells produce more BFU-E colonies (128±15 and 185±12 respectively) than CFUG/M/GM colonies (54±3 and 48±3 respectively). The number of multilineage CFU-GEMM colonies was the highest within 34DPACE ^+^ cells (16±5) and decreased two-folds in 34HCACE ^+^ cells (9±2) (Figure 3D). 34HCACE^-^ cell population did not produce CFU-GEMM (Figure 3D). Lineage distribution within colonies was confirmed by flow cytometry analysis of whole colonies obtained from each cell population. Indeed, the CD45 to Glycophorin-A-expressing cell ratio decreased from 34DPACE ^+^ cells to 34HCACE ^+^ and 34HCACE^-^ cell populations (97/3, 77/23 and 36/64 respectively) (Figure 3E). Flow cytometry analysis also demonstrated the presence of CD34^high^ immature progenitors in day 12 colonies for 34DPACE ^+^ and 34HCACE ^+^ cell population (2.09±0,48 and 0.68±0,14 respectively) whereas no CD34^high^cells were found in colonies derived from 34HCACE^-^cells (data not shown).

Altogether, our CFC-assays indicated that not only the progenitors contained in the 34DPACE ^+^ fraction presented a greater proliferative capacity but also had a higher multilineage potential than those contained in the 34HCACE ^+^ and 34HCACE^-^ EL fractions.

#### Long term culture-initiating cell assay on MS-5 stroma

We investigated the long-term hematopoietic potential of the different EL cell populations by seeding them on MS-5 stromal cells. 34DPACE^+^ and 34HCACE^+^ cells both developed round, stroma-adherent, HCs as well as cobblestone-like hematopoietic colonies underneath the stroma. 34HCACE^-^ cell cultures only produced round, stroma-adherent HCs and were devoid of cobblestone-like hematopoietic colonies. No HC production was observed from 34LDPACE^-^ and 34LHCACE^-^ cells (Figure 4A). Although 34DPACE^+^ and 34HCACE^+^ cells generated the same types of colonies, less cobblestone-like colonies were observed with 34HCACE^+^ cells compared to 34DPACE^+^ cells (data not shown). The kinetics of 34DPACE^+^, 34HCACE^+^ and 34HCACE^-^ hematopoietic productions were also remarkably different. While HC colonies were observed from day 3-4 of culture incipience with 34HCACE^+^ and 34HCACE^-^ cells, they were detected only from day 7-8 with 34DPACE^+^ cells. In this latter culture, colonies were observed beyond the 5 weeks of the standard long-term culture-initiating cell assay while no colony was found after three weeks with the 34HCACE^+^ and 34HCACE^-^ cells revealing a faster exhaustion. We next evaluated the kinetics of hematopoietic differentiation in 34DPACE^+^, 34HCACE^+^ and 34HCACE^-^ cells. Bulk HC cultures on MS-5 stroma were harvested at day 25 and 35 and stained with anti-CD45, -CD34 and -lineage-specific antibodies. Flow cytometry analysis indicated that for the 34DPACE^+^, 34HCACE^+^ and 34HCACE^-^ cell populations, all the cells expressed the pan-leukocyte marker CD45 and differentiated into myeloid (CD33^+,^ CD15^+^, CD14^+^), natural killer (CD94^+^) and B-lymphoid (CD19^+^) lineages (Figure 4B). However, loss of CD34^+^ progenitors and production of differentiated CD45^high^ cells were extremely different in the three cell populations. At 25 days of culture, only wells seeded with 34DPACE^+^ cells still contained a conspicuous level of CD34^+^ cells (8.5% of total CD34^+^ cells) that could be maintained over long term in the culture, whereas wells seeded with 34HCACE^+^ and 34HCACE^-^ contained only 2.37% and less than 1% of CD34^+^ cells respectively which disappeared after 35 weeks (Figure 4B). In addition, while undifferentiated CD45^low^ cells could still be observed after 35 days in the culture initiated with 34DPACE^+^ cells, 34HCACE^+^ and 34HCACE^-^ gave rise more rapidly to differentiated CD45^high^ cells (Figure 4B). Reanalysis of VEC and ACE at different time points of both cultures showed that both markers that typified newborn embryonic HSPCs, were quickly lost in culture, substantiating a rapid hematopoietic commitment of nascent HSPC on MS-5 (data not shown). The kinetics of lineage differentiation was also extremely different in the three populations. Although cells from the three populations underwent a series of differentiation steps leading first to the generation of myeloid cells and later on of B-lymphoid and NK cells, cultures seeded with 34DPACE^+^ cells presented a delayed but prolonged production of all lineages compared to cultures initiated with 34HCACE^+^ and 34HCACE^-^ cells (Figure 4B). Indeed, after 25 days and even after 35 days, more myeloid cells were present in the culture initiated with 34DPACE^+^ cells than with 34HCACE^+^ and 34HCACE^-^ cells. Of note, 34HCACE^+^ and 34HCACE^-^ cells have already produced myeloid cells at earlier time points of culture. 34DPACE^+^ cells also generated B-lymphoid and NK cells with a delay compared to 34HCACE^+^ and 34HCACE^-^ cells (Figure 4B).

**Figure 4.**
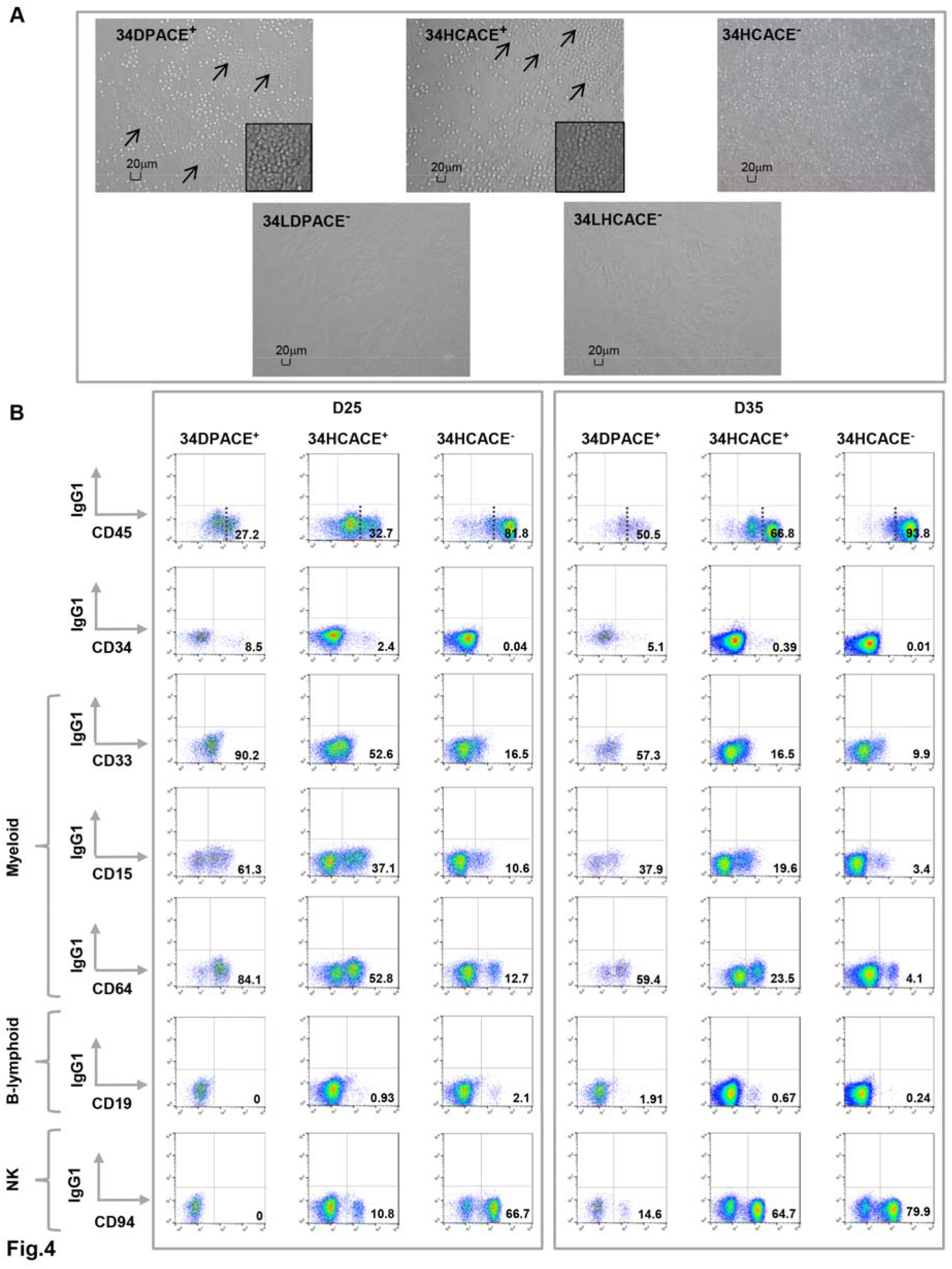
Long-term hematopoietic potential of the 34DPACE^+^, 34HCACE^+^ and 34HCACE^-^ EL cell fractions. **(A)** Typical packed round, stroma-adherent HCs for 34DPACE^+^, 34HCACE^+^ and 34HCACE^-^ cell populations and phase-dark cobblestone-shaped colonies underneath the stroma for 34DPACE^+^ and 34HCACE^+^ cell populations after 25 days of culture on MS-5. No hematopoietic colony was observed on MS-5 for 34LDPACE^-^ and 34LHCACE^-^ cell population. Arrows pointed to phase-dark cobblestone-shaped colonies. Smaller rectangles show enlargements of phase-dark cobblestone-shaped colonies with 1,5 fold magnification. Scale bar = 20μm. **(B)** CD45, CD34 and lineage-specific stainings of HCs obtained after 25 and 35 days of culture on MS-55 for 34DPACE^+^, 34HCACE^+^ and 34HCACE^-^ EL populations. Numbers indicate the percentages of positive cells for each marker. For CD45 dot plot, dotted lines discriminate between CD45^low^ and CD45^high^ cells and percentage correspond to CD45^high^ cells. **(A-B)**: Data are representative of at least three independent experiments performed on 6.2 to 8.4-week-old ELs.

In conclusion, our long-term culture assay, demonstrated that 34DPACE^+^ cells showed a delayed but prolonged hematopoietic production when cultured on MS-5 cells, with a sustained maintenance of undifferentiated CD34^+^ CD45^low^ cells compared to 34HCACE^+^ and 34HCACE^-^ EL cells.

#### CSFE analysis

The proliferative capacity of the different EL cell subsets was probed at day 21 and 35 of MS-5 co-cultures using CFSE staining. At 21 days of culture, 34DPACE^+^ -derived CD45^+^ cells displayed more divisions than 34HCACE^+^ - and 34HCACE^-^-derived CD45^+^ counterparts. Of note, as early as day 3 after CSFE labeling, 34DPACE^+^ cells already produced at least one or two more progenies than 34HCACE^+^ and 34HCACE^-^ cells (Figure 5A). Moreover, the first peak of division for 34DPACE^+^ contained only 25.58% of cells whereas it still contained 47.2% and 52.67% of cells for 34HCACE^+^ and 34HCACE^-^ cultures respectively (Figure 5A). At day 6 after CSFE labeling, most 34DPACE^+^ cells had performed up to 6 divisions and the first peak of proliferation was only containing 4.71% of cells while 34HCACE^+^ and 34HCACE^-^ cells only produced 5 successive generations of HCs and the first peak of proliferation was still containing 15.32% and 26.85% of cells respectively (Figure 5A). At 35 day of culture, 34DPACE^+^ cells generated the same number of progeny than the 34HCACE^+^ and 34HCACE^-^ cultures (Figure 5B) but produced more HCs with high proliferative potential than for 34HCACE^+^ and 34HCACE^-^ cells. Indeed, after 3 and 6 days of CSFE labeling the first peak of proliferation only contain 39.66% and 9.95% of cells for 34DPACE^+^ co-culture whereas it still contain 56.98% and 43% and 70.85% and 53.28% of cells for 34HCACE^+^ and 34HCACE^-^ cultures respectively (Figure 5B) hence highlighting the superior proliferative response of 34DPACE^+^ cells when co-cultured on MS-5.

**Figure 5.**
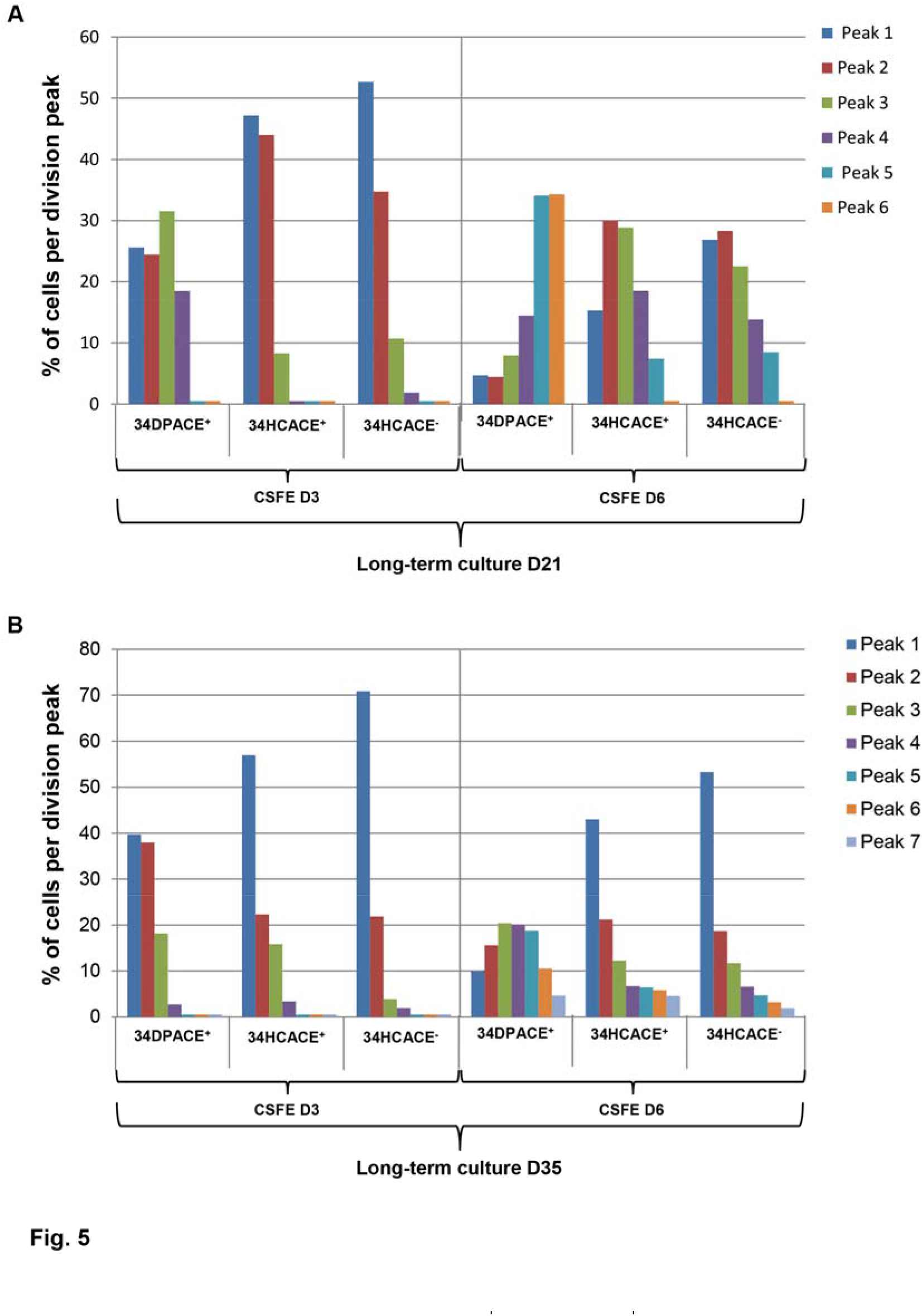
Proliferative potential of 34DPACE^+^, 34HCACE^+^ and 34HCACE^-^ EL cell fractions in long-term culture assays. (**A-B**) Kinetics of proliferation. HC derived from each EL cell population were stained with CSFE after 21 (**A**) and 35 days (**B**) of culture on MS-5. CFSE-labeled HC were cultivated for 3 and 6 additional days. The percentage of proliferating cells was analyzed by flow cytometry based on CFSE dilution at day 3 and 6 after staining with an anti-CD45 Mab. Each histogram bar represents the percent of cell per division peak in the CD45 cell fraction for each cell population analyzed. Peak 1 corresponds to the number of cell divisions between day 0 undivided cells and the first peak obtained at day 3 and day 6. Following peaks represent one or more additional division based on their fluorescence intensity. **(A-B):** Data are representative of 3 independent experiments performed on 6.4 to 7.7-week-old ELs. D: day. CFSE, 5,6-carboxyfluorescein-diacetate-succinimidyl-ester.

#### Cell cycle analysis

We also performed cell cycle experiments using either Hoechst alone or both Hoechst and Pyronin Y. Hoechst discriminated between cells in G_0_/G_1_ and cells in S/G_2_/M. Most of the 34DPACE^+^ cells were cycling (83.5% of S/G_2_/M) while only 33.1% and 10.5% of the 34HCACE^+^ and 34HCACE^-^ cells were in a similar state respectively (Figure 6A). Pyronin Y/Hoechst allowed separating cells in G_0_ from those in G_1_. The proportion of quiescent i.e. G_0_, cells was only 3.34% in the 34DPACE^+^ population while it reached 31.9% and 27.3% in the 34HCACE^+^ and 34HCACE^-^ cell populations respectively (Figure 6B). Thus, cells switched abruptly from an actively dividing to a quiescent state when 34DPACE^+^ cells were considered and compared to 34HCACE^+^ and 34HCACE^-^cell populations. Cell cycle analysis combined with *ex vivo* assay indicated a higher expansion capacity of 34DPACE^+^ cell subset compared to 34HCACE^+^ and 34HCACE^-^ subsets likely due to a better capacity to proliferate and to undergo self-renewal to maintain/expand undifferentiated HSPCs *ex vivo*.

**Figure 6.**
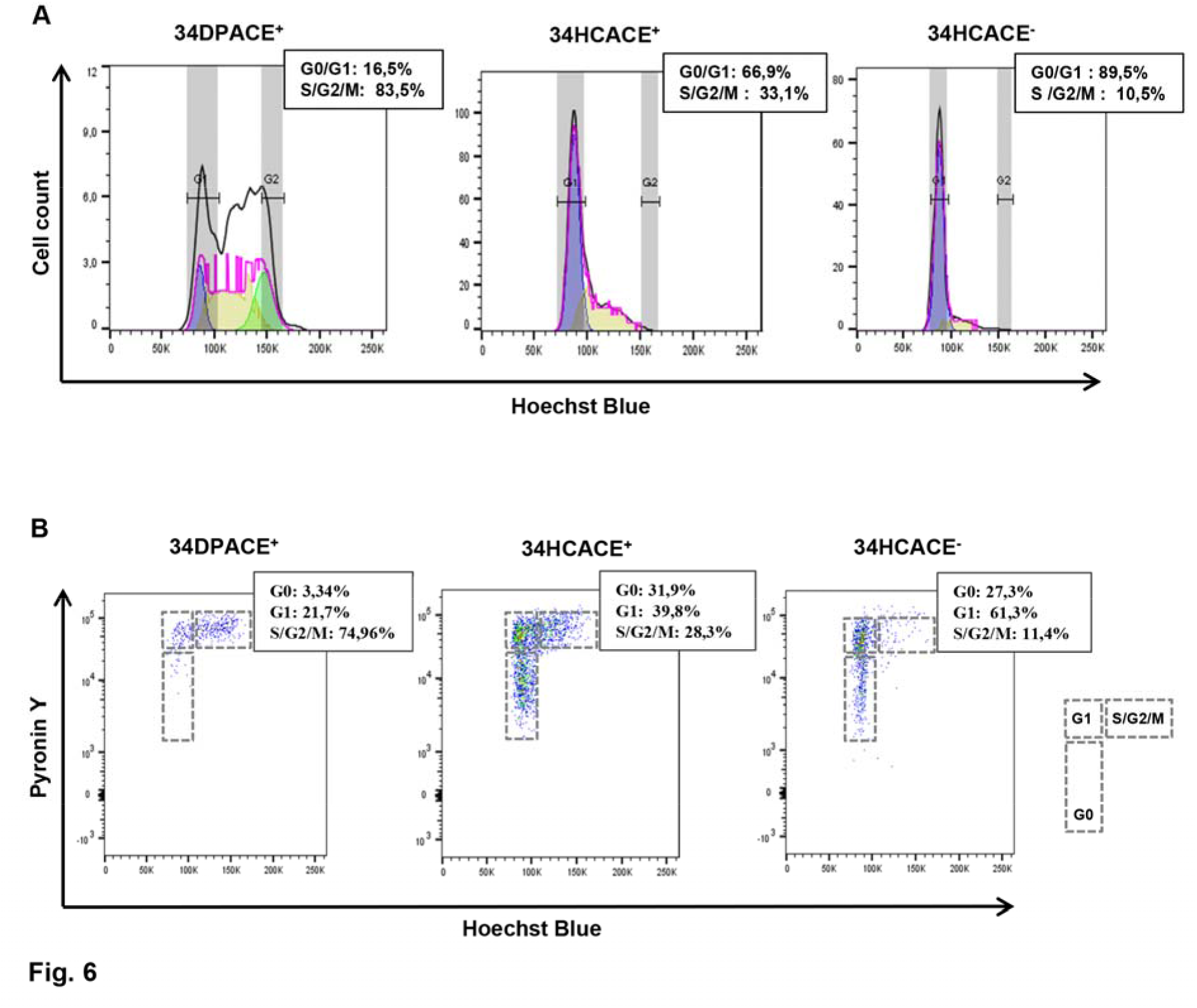
Cell cycle status of 34DPACE^+^, 34HCACE^+^ and 34HCACE^-^ EL cell fractions. **(A-B)** Whole liver cells from a 7.7-week-old human embryo stained with Hoechst and anti-CD45, -VEC, -ACE- and -CD34 Mabs or with Hoechst, Pyronin Y and anti-CD45, -VEC, -ACE- and -CD34 Mabs. **(A)** Cell cycle analysis based on Hoechst fluorescence intensity analyzed on each gated populations. Representative FACS histograms for 34DPACE^+^ (left), 34HCACE^+^ (middle) and 34HCACE^-^ (right) EL cell populations. The percentage of cells in the G_0_/G_1_ and S/G_2_/M phases within the different populations is indicated on the top of each histogram. **(B)** Cell cycle analysis based on Hoechst/Pyronin Y fluorescence intensity analyzed on each gated population. Representative FACS dot plot for 34DPACE^+^ (left), 34HCACE^+^ (middle) and 34HCACE^-^(right) EL cell populations. The percentage of cells in the G_0_, G_1_ and S/G_2_/M phases within the different populations is shown on the top of each dot plot. **(A-B):** Histogram and dot plots show the results from 1 representative experiment (n = 3) performed on 6.8 to 8.2 week-old ELs.

### *In Vivo* Long term reconstituting ability is restricted to the 34DPACE^+^ EL subset

To determine which cell population present in the human EL possessed the highest ability to engraft into immunocompromised mice, EL cell subsets were transplanted into sub-lethally irradiated NSG mice. As a positive control we used the CD34^+^ CD45^+^ HSPC population that included the 34DPACE^+^, 34HCACE^+^ and 34HCACE^-^ cell fractions. Engraftment was based on flow cytometry analysis of murine BM cells harvested 4 months after transplantation and was considered positive when more than 0.1% human CD45^+^ cells were detected.

Different doses ranging from 1000 to 20,000 34DPACE^+^ cells per mouse were injected. A dose-dependent increase was observed. Two out of 2 mice transplanted with as few as 5000 cells exhibited positive engraftment at 4 months. When 10.000 and 20.000 cells were injected, 5 out of 5 mice exhibited positive engraftment. The percent of repopulating cells in the murine BM reached 87.5% when 20,000 cells were injected (Supplemental Table S4). Conversely, no engraftment was observed in mice transplanted with 5000 34HCACE^+^ cells (Supplemental Table S4) and very few human cells (0.2%) were detected in one out of 6 mice engrafted with 10,000 34HCACE^+^ cells. Low reconstitution (0.3%) could be observed in one mouse out of 6 when 100,000 34HCACE^-^ cells were injected. One out of 2 mice injected with 50,000 CD34^+^ CD45^+^ control cells exhibited a significant level of BM engraftment (70.5%) but when less than 50,000 CD45^+^ CD34^+^ cells were injected (4 mice tested), no engraftment was detected (Supplemental Table S4).

We then analyzed the phenotype of human repopulating cells with a panel of human-specific antibodies including CD34, CD38, CD45, myeloid (CD33, CD15, CD14), B-lymphoid (CD19) and T- lymphoid (CD3) lineage-specific antibodies. Multilineage reconstitution with the typical proportional predominance of human B-lymphoid cells (about 65% of CD19^+^ B-lymphocytes, 25% of CD33^+^ myeloid cells and less than 1% of CD3^+^ T-lymphoid cells) was observed in the BM of mice that received 34DPACE^+^ cells or CD45^+^ CD34^+^ control cells at 3 months post-transplantation (Figure 7A). The same proportion of human cell subsets was found when mice were reconstituted with 34HCACE^+^ and 34HCACE^-^ cells (data not shown). Additionally, a significant fraction of primitive CD34^+^ CD38^low^ (3.1% and 3.7% respectively) human progenitors were still found in the BM of mice injected with 34DPACE^+^ cells or CD34^+^ CD45^+^ control cells (Figure 7A).

**Figure 7:**
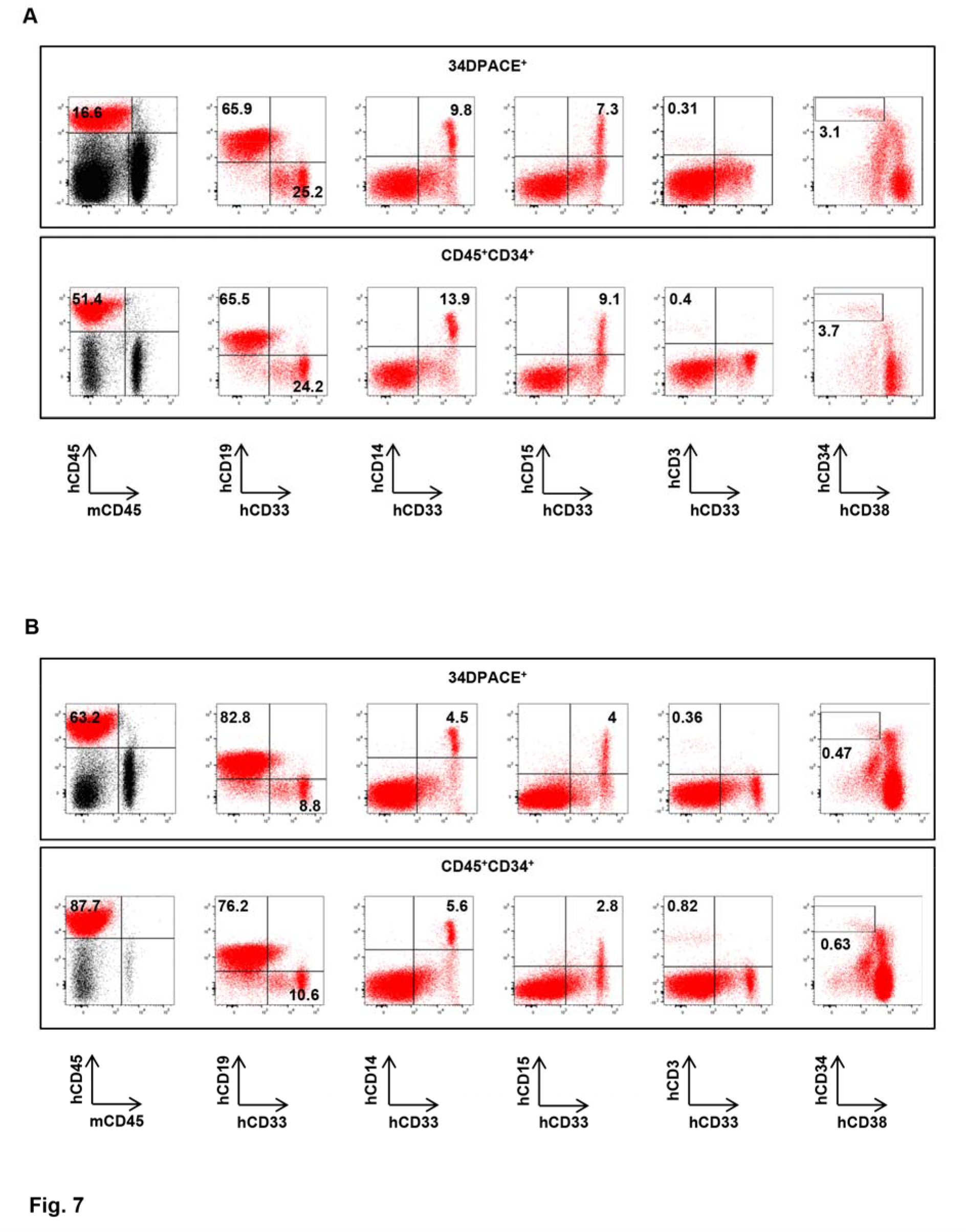
Primary and secondary hematopoietic reconstitution of NSG mice by 34DPACE^+^ compared to CD45^+^ CD34^+^ EL cell fractions. **(A)** 5000 34DPACE^+^ and 50.000 CD45^+^ CD34^+^ cells sorted from a 7.2-week-old human EL cells injected into sublethally irradiated primary NSG mice. Primary transplanted mice were tested for the presence of human cells in their BM four months after transplantation. Representative flow cytometry analysis for 34DPACE^+^ and CD34^+^ CD45^+^ cell populations demonstrated multilineage engraftment with a typical proportion of B-lymphoid and myeloid cells as assessed by multiple-staining with human specific Mabs to CD45, myeloid (CD33, CD14, CD15), B-lymphoid (CD19) and T-lymphoid (CD3) cells. Additionally, a significant proportion of primitive CD34^+^ CD38^low^ cells were still found in the BM of mice injected with 34DPACE^+^ and CD34^+^ CD45^+^ cells as assessed by triple-staining with human specific Mabs to CD45, CD34 and CD38 cells. **(B)** 3.10^6^ human cells sorted from the BM of primary engrafted mice were injected into secondary sublethally irradiated NSG mice. Secondary transplanted mice were tested for the presence of human cells in their BM four months after transplantation. Representative flow cytometry analysis for 34DPACE^+^ and CD34^+^ CD45^+^ cells demonstrated multilineage engraftment with a typical proportion of B-lymphoid and myeloid cells as assessed by multiple-staining with human specific Mabs to CD45, myeloid (CD33, CD14, CD15), B-lymphoid (CD19) and T-lymphoid (CD3) cells. Additionally, a significant proportion of primitive CD34^+^ CD38^low^ cells were still found in the BM of mice injected with 34DPACE^+^ and CD34^+^ CD45^+^ cells as assessed by triple-staining with human specific Mabs to CD45, CD34 and CD38 cells. **(A-B):** A murine CD45 Mab was included in all stainings to check the specificity of human Mabs. Numbers indicate the percentages of positive cells in the corresponding quadrants. Data are representative of 6 independent experiments performed on 7.2 to 8.2-week-old ELs.

We performed secondary reconstitutions with human cells isolated from the BM of the primary mice reconstituted with the 34DPACE^+^ and CD34^+^ CD45^+^ populations. 3.10^6^ human cells isolated from mice that received 5000 34DPACE^+^ cells were injected into three secondary recipients. The same design of experiments was applied to mice that received 50,000 CD34^+^ CD45^+^ cells as control. All secondary recipients transplanted with either 34DPACE^+^ or CD34^+^ CD45^+^ exhibited high levels of human cells in their BM 3 months post-transplantation (Supplemental Table S5). Secondary recipients transplanted with either 34DPACE^+^ cells or CD34^+^ CD45^+^ cells demonstrated multilineage engraftment with Blymphocytes (81.8 and 76.2% respectively) and myeloid cells (8.8 and 10.6% respectively) as well as few T-lymphocytes (0.66 and 0.82% respectively) (Figure 7B). As for primary reconstitution a significant fraction of primitive CD34^+^ CD38^low^ (0.47% and 1.64% respectively) human progenitors were still found in the BM of mice injected with 34DPACE^+^ or CD34^+^ CD45^+^ control cells (Figure 7B). These results further indicated that long term repopulating potential was restricted to the 34DPACE^+^ cell subset in the human EL.

### Absence of a VEC^low^CD45^low^ DP population in neonatal CB and adult BM

To determine whether 34DP cells were still present in neonatal and adult life, we flow cytometry analyzed term CB and adult BM cells. No VEC^low^CD45^low^ DP cells could be found in mononuclear cells prepared from term CB (Supplemental Figure S3). We were only able to detect VEC^-^CD45^low^ and VEC^-^CD45^high^ HCs and very few VEC^+^ CD45^-^ ECs (Supplemental Figure S3). A rare population (0.11%) of cells co-expressing VEC and CD45 could be detected in more differentiated CD45^high^cells (Supplemental Figure S3). These latter cells were mostly CD34^-^ and co-expressed myeloid lineage markers (data not shown).

FACS profiles obtained from BM mononuclear cells were identical to that obtained from CB ones except that we could detect more VEC^+^ CD45^-^ECs (Supplemental Figure S3).

### Identification of a VEC^low^CD45^low^ DP population in human embryonic YS, AGM and placenta

We investigated whether earlier hematopoietic sites of the developing human conceptus also harbored cells co-expressing VEC and CD45. We thus performed flow cytometry analyzes on human YS, AGM and placenta (Supplemental Table S6). In YS and AGM, although VEC and CD45 expression were largely mutually exclusive, a rare, double positive population (0.047 + /-0.023 and 0.045 + /-0.028% respectively) was detected (Figure 8A and Supplemental Table S6) as previously described in the EL (Oberlin et al., 2010) and herein. In addition, VEC^**+**^ CD45^-^ endothelial and VEC^-^CD45^+^ hematopoietic subsets were also identified (Figure 8A). Frequency of the DP subset in the YS and AGM changed along ontogeny with the highest proportion observed at the earliest stages analyzed (Supplemental Table S6). No DP cells were observed in AGM and YS dissected from embryos older than 6.6 week-old (Supplemental Table S6). Additional flow-cytometry analysis of CD34 and ACE in combination with VEC and CD45 demonstrated that DP cell-coexpressing VEC and CD45 in the YS and AGM were mainly positive for CD34 (Figure 8B) and ACE (data not shown).

**Figure 8.**
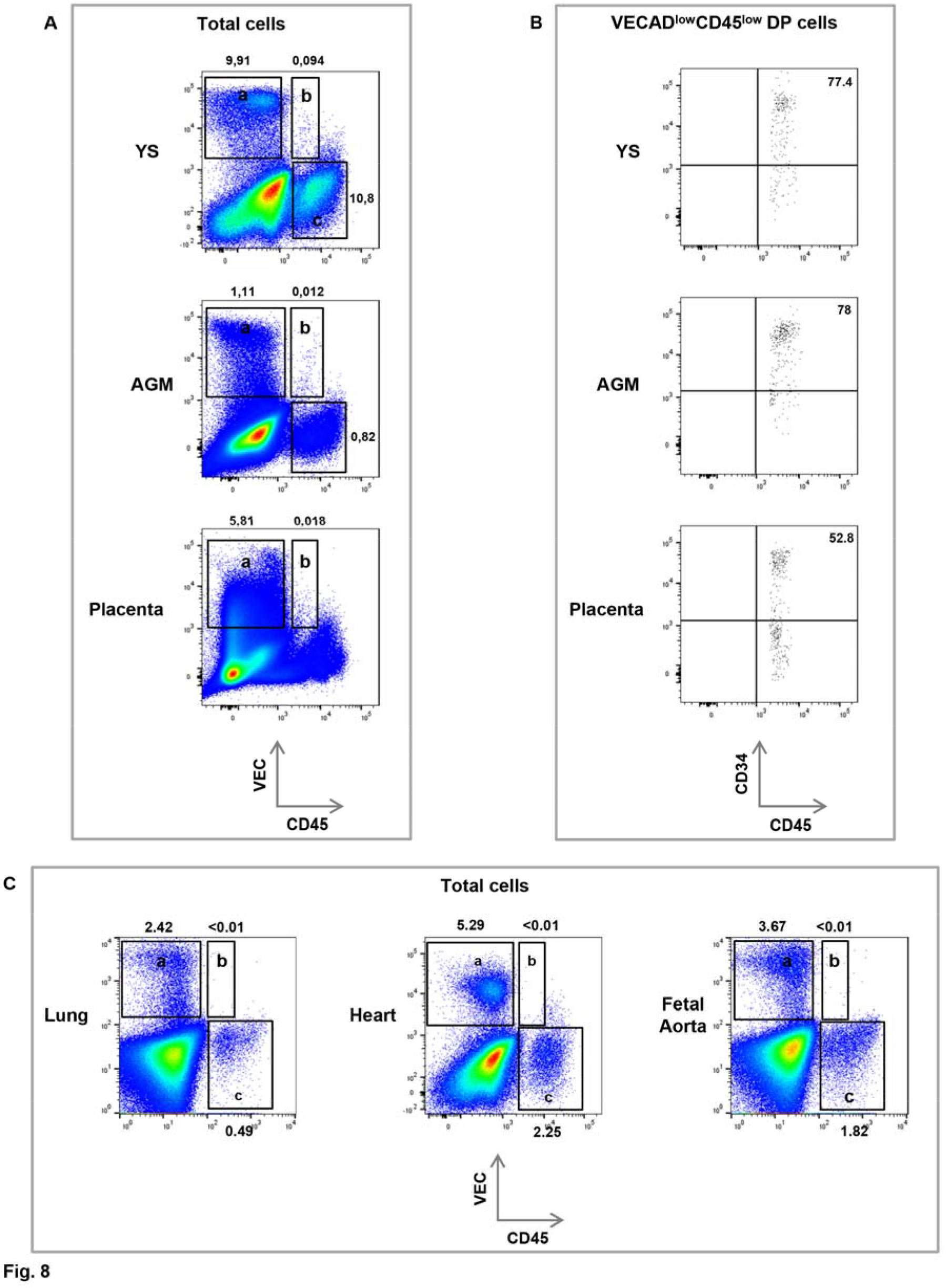
Identification of EC, DP and HC fractions in human embryonic and fetal tissues. (**A-B**) Whole YS, AGM and placenta cells from a 6.3-week-old human embryo, were triple-stained with anti-CD45, -VEC and -CD34 Mabs. **(A)** Flow cytometry analysis demonstrating endothelial (VEC^+^ CD45^-^ -frame **a**-), DP (VEC^low^CD45^low^ -frame **b**-) and hematopoietic (VEC^-^CD45^+^ -frame **c**-) cell populations. **(B)** Flow cytometry analysis showing CD34 cell surface antigen expression in the gated DP (VEC^low^CD45^low^) cell population. **(A-B):** Numbers indicate the percentages of positive cells in the corresponding frames. Data are representative of at least three independent experiments performed on 5 to 6.3-week-old embryos. **(C)** Whole aorta, lung and heart from a 7.4-week-old human embryo triple-stained with anti-CD45, -VE-cadherin and -CD34 Mabs. Flow cytometry analysis showing endothelial (VECAD^+^ CD45^-^ -frame **a**-) and hematopoietic (VECAD^-^CD45^+^ -frame **c**-) cell populations but not DP (VECAD^low^CD45^low^ -frame **b**-) cell population. **(C):** Data are representative of at least three independent experiments performed on 5 to 9.6-week-old embryos.

In contrast, 5 to 6.3 week placenta showed only few DP cells (0.015 + /-0.0033) even at the earliest stages analyzed (Figure 8A and Supplemental Table S6). No DP cells were observed in placenta dissected from embryos older than 6.6 week-old. As in the AGM and YS, endothelial and hematopoietic subsets were also identified (Figure 8A). Flow-cytometry analysis of CD34 in combination with VEC and CD45 demonstrated that DP cell-coexpressing VEC and CD45 in the placenta were also mainly positive for CD34 but to a lesser extend compared to the YS and the AGM (Figure 8B). In all stages tested, umbilical cord, heart, lung, head and fetal aorta (considered as nonhematopoietic organs) did not contain (<0,01%) DP cells while endothelial and hematopoietic subsets were identified (Figure 8C and Supplemental Table S6).

### Extended long term hematopoietic capacity of the 34DP population in the early human embryo

We tested whether the 34DP (VEC^low^CD45^low^CD34^+^) population isolated from the early embryo YS, AGM and placenta was endowed with hematopoietic potential. The long-term hematopoietic potential, probed on MS-5 stromal cells, of 34DP cells sorted from YS, AGM and placenta was compared to that of 34HCs (VEC^-^CD45^low^CD34^+^) sorted from the same organs or sites. 34DP cells and 34HCs sorted from EL were also seeded on MS-5 cells for comparison. As for the EL, both 34DP cells and 34HC cells sorted from the YS and AGM developed round, stroma-adherent HC colonies as well as cobblestone-like HC colonies underneath the stroma (Figure 9A). While hematopoietic colonies were observed as soon as 3-4 days in the cultures initiated with 34HCs, they were detected only after 7-8 days in cultures initiated with 34DP cells. Conversely extended-hematopoietic activity, scored beyond the 5 weeks of the standard long-term culture assay, was detected exclusively within the 34DP cell population. No hematopoietic activity was observed from 34DP cells sorted from the placenta probably due to the low amount of cells seeded. Hematopoietic activity was detected from 34HCs (data not shown) but remained restricted below 5 weeks of culture.

**Figure 9.**
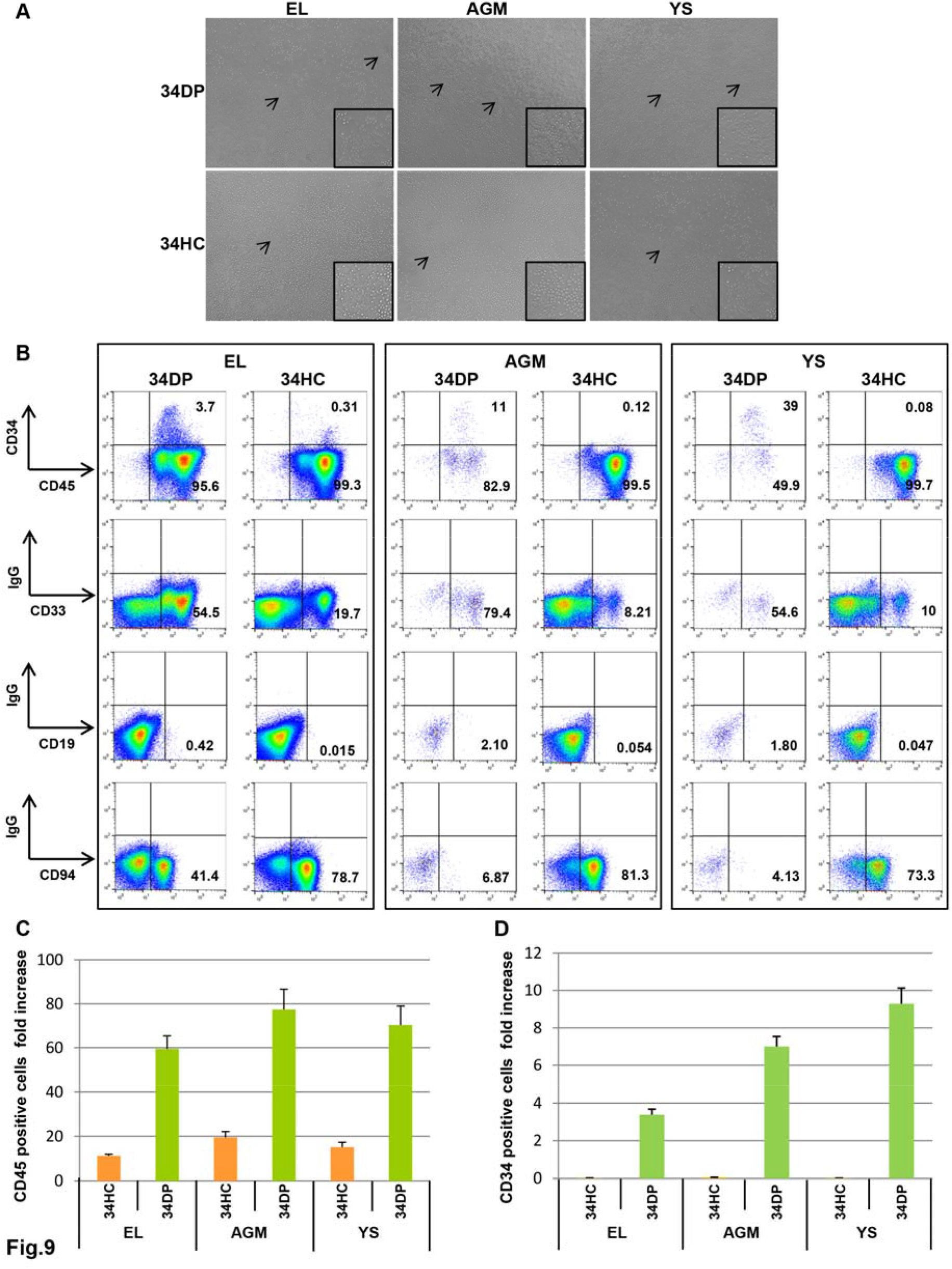
Long-term hematopoietic potential of 34DP (VEC^low^CD45^low^CD34^high^) and 34HC (VEC^-^CD45^low^CD34^high^) EL, AGM and YS cell fractions. 34DP and 34HC sorted from EL, AGM and YS cells were grown on MS-5 stromal cells in bulk (**A-E**) **(A)** Typical colonies of packed round, stroma-adherent HCs and phase-dark cobblestone-shaped colonies underneath the stroma obtained after 25 days of culture on MS-5 for 34DP and 34HC EL, AGM and YS cell populations. Arrows pointed to phase-dark cobblestone-shaped colonies. Frames show enlargement of, phase-dark cobblestone-shaped, colonies with 1,5 fold magnification. Scale bar = 20μm. (**B**) CD34^+^ CD45^low^ HSPC and myeloid (CD33), NK (CD94) and B-lymphoid (CD19) cells production by 34DP and 34HC EL, AGM and YS cells after 25 days of culture on MS-5 cells. Numbers indicate percentages of positive cells in the corresponding quadrants. **(A-B):** Data are representative of three independent experiments performed on 5 to 6.2-week-old embryos. **(C)** CD45^+^ cell number fold increase for 34DP and 34HC EL, AGM and YS cell populations after 25 days of culture on MS-5. **(D)** CD34^+^ cell number fold increase for 34DP and 34HC El, AGM and YS cell populations after 25 days of culture on MS-5. **(C-D)**: Each histogram represents the mean value + /- standard deviation of three independent experiments performed on 5 to 6.2-week-old embryos.

The kinetic of hematopoietic differentiation of 34DP cells compared to 34HCs from the same organ were also compared. At 25 days of co-culture 34DP and 34HC population progenies both expressed the pan-leukocyte marker CD45 and displayed myeloid (CD33^+^), natural killer (CD94^+^) and Blymphoid (CD19^+^) lineage differentiation (Figure 9B). However, loss of undifferentiated CD45^low^CD34^+^ hematopoietic progenitors was extremely different in both cultures. Indeed, after 25 days of culture, only wells seeded with 34DP cells still contained high level of CD45^low^CD34^+^ cells (3.7%, 11%, 39% for the EL, the AGM and the YS respectively), whereas wells seeded with 34HCs contained only 0.31, 0.12 and 0.08% of CD45^low^CD34^+^ cells for the EL, the AGM and the YS respectively (Figure 9B). The kinetic of lineage differentiation was also extremely different in both cultures. Although cells from both populations underwent a series of differentiation steps leading first to the generation of myeloid cells and later on to B-lymphoid and NK cells, all the cultures seeded with 34DP cells presented a delayed production of all lineages compared to cultures initiated with 34HCs (Figure 9B). To compare the proliferative capacity of 34DP to 34HC cells, the total progeny produced from each cell population was measured after 25 days of culture. As shown on Figure 9C the magnitude of amplification of CD45^+^ cells for 34DP cells was 5.3, 4.1 and 4.7 fold higher than for 34HCs in the EL, the AGM and the YS respectively. We also measured the CD34^+^ cell production and demonstrated that the magnitude of amplification of CD34^+^ cells was of 3.4±0.30, 7.1±0.54 and 9.3±0.83 for 34DP cells in the EL, the AGM and the YS respectively, whereas in culture initiated with 34HCs no CD34 amplification could be observed (Figure 9D).

Altogether, long-term culture assays clearly showed that similar to the EL, 34DP cells sorted from YS and AGM were able to maintain undifferentiated CD45^low^CD34^+^ progenitors for a longer period of time than 34HCs, and to yield all blood cell progenies with a delayed but prolonged kinetics of hematopoietic differentiation compared to 34HCs.

## DISCUSSION

Identification of the different hematopoietic populations operating in the early human embryo and their hierarchical organization is of paramount importance to decipher the cellular and molecular mechanisms associated with self-renewal and differentiation and to design innovative cell based therapies.

In order to distinguish more accurately the earliest HSPCs emerging in the human embryo, we performed here a complete ensemble of phenotypic and functional characterizations of the 34DP and the 34HC EL cells using markers commonly used to identify human HSPC and endothelial cells and identified ACE as a key marker that, associated with VEC, allows discriminating hematopoietic populations endowed with a strong hematopoietic potential from those with less potential.

### Identification of 34DPACE^+^ cells at the top of the hierarchy in the human EL

Our extensive phenotypic analysis extended the phenotype of the DP cells (Oberlin et al., 2010a) to CD34^+^ VEC^+^ CD105^+^ CD90^+^ CD117^+^ CD45RA^-^CD38^-/low^ CD45^+^ cells confirming their hematoendothelial phenotype and their enrichment in HSPC markers compared to HCs.

We then demonstrated that ACE was an extremely helpful marker to refine the EL HSCP hierarchy. Functional analysis demonstrated that VEC, CD45 and CD34 in conjunction with ACE allow separating three subpopulations endowed with contrasted hematopoietic potential. Although all three 34DPACE^+^, 34HCACE^+^ and 34HCACE^-^ cell populations possess multilineage differentiation potential, only the former could maintain/expand undifferentiated HSPCs and robustly expand *in vitro* and *in vivo*. This places the 34DPACE^+^ population at the top of the EL hematopoietic hierarchy. ACE, a regulator of blood pressure, is a cell surface marker of adult HSPCs (Jokubaitis et al 2008). It is expressed in all developing blood-forming tissues of the human embryo and fetus (Sinka et al 2012) and has been reported as expressed by the intra-aortic hematopoietic clusters of the AGM, as well as by surrounding ECs (Sinka et al., 2012). In keeping with this, we demonstrated that ACE is tightly associated with EC and DP cells but not with more mature HCs in the EL. Our results are also in agreement with the fact that human fetal liver derived CD34^+^ ACE^+^ cells, but not CD34^+^ ACE^-^ cells, are endowed with LTC-IC potential and sustain multilineage hematopoietic cell engraftment when transplanted into NOD/SCID mice (Sinka et al 2012). However when splitting EL HSPC into 34DPACE^+^, 34HCACE^+^ and 34HCACE^-^ cell population we observe a radical change from a more powerful VEC^+^ subpopulation to less powerful VEC^-^ subsets. For that reason we place the 34DPACE^+^ at the top of the EL hematopoietic hierarchy. Since VEC and ACE are progressively lost with time as liver development proceeds, 34DPACE^+^ could thus represent a transient HSPC population harboring endothelial traits and originating from earlier hematopoietic sites. This population will, supposedly, progressively give rise first to 34HCACE + and then to 34HCACE^-^ cells following the subsequent loss of VEC and ACE (See recapitulative scheme in Figure 10).

**Figure 10.**
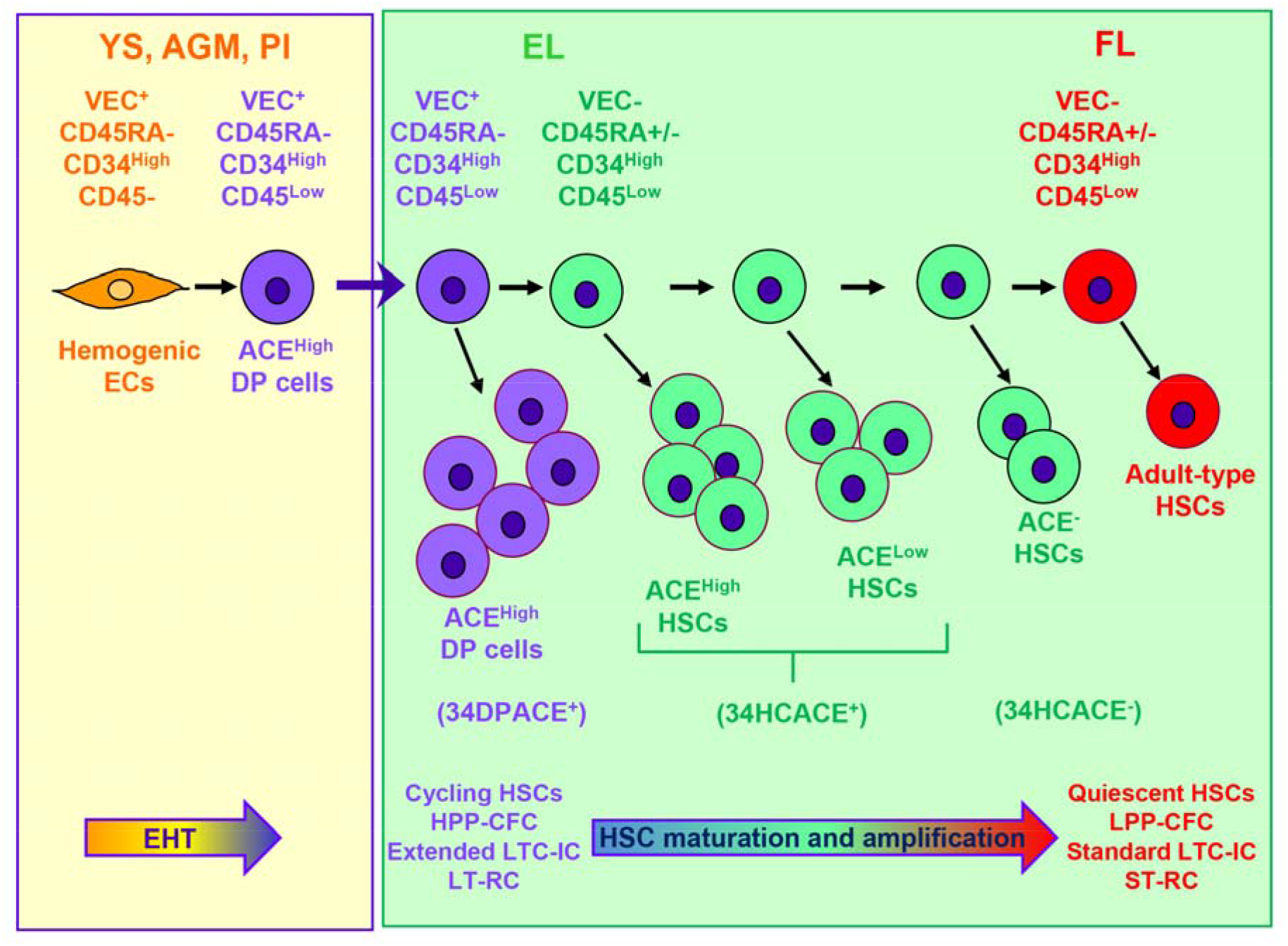
HSPC hierarchy in the human embryo. During human embryonic development different waves of hemogenic endothelial cells (ECs) are produced in the yolk sac (YS), AGM and potentially in the placenta (Pl). These hemogenic ECs give rise to CD45^+^ CD34^+^ VEC^+^ ACE^+^ DP cells that colonize the embryonic liver (EL). Quickly after EL colonization, DP cells loose VE-cadherin, then ACE. Their progeny have reduced self-renewal capacity and contain lower LTC-IC as well as *in vivo* repopulating cells compared to their parent cells. After several cell division and maturation steps embryonic HSPC will give rise to *bona fide* adult HSPCs. EHT, Endo-Hematopoietic Transition; LPP-CFC, Low Proliferative Potential-Colony Forming Cells; HPP-CFC, High Proliferative Potential-Colony Forming Cells; LTC-IC, Long Term Culture-Initiating Cells; LT-RC, Long Term-Repopulating Cells; ST-RC, Short Term Repopulating Cells; FL, Fetal Liver.

Cell cycle analysis also demonstrated that cells switched abruptly from an actively dividing to a quiescent state when 34DPACE^+^, 34HCACE^+^ and 34HCACE^-^ cell populations are considered respectively. Unlike adult bone marrow HSPCs which are mostly quiescent, fetal liver HSPCs are thought to be highly proliferative (Bowie et al., 2007, 2006; Morrison et al., 1995), leading to a ∼38-fold expansion from E12 to E16 in the mouse (Ema and Nakauchi, 2000). Here we show that, unexpectedly, the proliferative potential is unequally distributed among the different EL hematopoietic populations. A contrasted proliferation potential has already been demonstrated in the murine AGM, using Fucci reporter mice to visualize the cell-cycle status. A slowing down of cell cycle is observed as cells start to acquire a definitive HSC state immediately before reaching the fetal liver, similar as we report for EL HSCs (Batsivari et al., 2017). Whether or not this multiplication landscape is conserved when cells reached the fetal liver is yet to be defined. Comparison of the EL hematopoietic cell populations to that of quiescent adult bone marrow HSPC (Arai and Suda, 2007; Wilson et al., 2008) will provide mechanistic insights into how to tailor HSPC expansion *ex vivo*.

Our phenotypic study combine to *in vitro* and *in vivo* hematopoietic analysis not only shed a new light on the hierarchical organization of the human EL HSPC compartment but also confirms that during EHT the hematopoietic lineage seems to be gradual rather than abrupt. Such a hierarchical hematopoietic organization has been described within intra-aortic hematopoietic clusters in the murine and human AGM (Boisset et al., 2014; Rybtsov et al., 2014 and 2011; Ivanovs et al., 2014) and in the murine fetal liver (Kieusseian et al., 2012). Hierarchical organization of the human EL remain largely unexplored at the exception of a recent study which describes a novel surface marker, glycosylphosphatidylinositol-anchored protein GPI-80, that is functionally required for HSPC self-renewal at fetal stages (Prashad et al., 2014). Although we have not addressed the biological value of GPI-80, it will be interesting to investigate whether differences in levels of GPI-80 expression in DP cells can still help to even refine EL HSCP hierarchy at embryonic stages.

### Identification of 34DP cells in the early human embryo but not in neonatal or adult hematopoietic tissues

Based on our previous studies in the EL (Oberlin et al., 2010a, b), we also addressed the question of the biological value of VEC and CD45 expression in the human YS and the AGM to isolate cells carrying a strong amplification and self-renewal potential. We do find DP cells, most of which are positive for CD34 and ACE in these sites. Moreover, the extended long-term hematopoietic capacity in the AGM and the YS is restricted to DP cells. These results are in agreement with studies in the mouse (Taoudi et al., 2005 and 2008, Fraser et al., 2003, North et al., 2002; Nishikawa et al., 1998a) and the human embryo (Ivanovs et al., 2014) showing that the first YS and AGM HSPCs harbored a dual hemato-endothelial phenotype. Furthermore, detection of DP cells as early as 5 weeks of gestation in the human EL and at the same time in the AGM region and YS, two sites of hematopoietic emergence, suggests that the EL is seeded by AGM or YS-derived DP cells.

In contrast to the YS and AGM, the 5 to 9.6 week placentas show only few DP cells. Despite the fact that CD34^+^ cells are detected as early as week 5 of gestation in the human placenta (Barcena et al., 2009), repopulating HSPCs are present at week 6 of gestation (Robin et al., 2009) although two studies report the presence of transplantable HSPCs only after week 9 of gestation or even later, as determined by xenotransplantation into immunocompromised mice (Ivanovs 2011; Muench et al., 2017). These latest observations and the fact that we could only identify few DP cells in the 5-9.6 week placenta highlight the possibility that, contrary to the mouse embryo (Rhodes et al., 2008), the human embryo placenta may not be a potent site of HSPC emergence. Regarding amplification, our observations are also different from those reported for the mouse placenta that is considered as a site of unbiased HSPC amplification (Gekas et al., 2005; Ottersbach and Dzierzak, 2005).

As expected, in all stages tested, non-hematopoietic organs as umbilical cord, heart, head and lung, only contain extremely few cells co-expressing VE-cadherin and CD45 (<0.1%). As for the placenta, these DP cells could represent embryonic HSPCs migrating from the YS and the AGM to the EL. Finally, VEC expression is absent from neonatal and adult hematopoietic tissues. This indicates that the DP fraction progressively loose VEC expression as development proceeds to completely disappear from neonatal stages.

### Role of VEC in human HSC emergence, migration and amplification

Self-renewal and multilineage reconstitution is tightly associated with VEC in the early human embryo. Does VEC represent a signature of endothelial origin that disappears with time or is VEC expression associated with hematopoietic-specific molecular pathways to sustain EL hematopoiesis? VEC is one of the classic Ca^2+^ dependent, homophilic adhesion molecules primarily expressed in endothelial cell adherens junctions (Lampugnani et al., 1992). The intracellular domain of VEC physically interacts with p120 catenin, beta-catenin, alpha-catenin, and the actin cytoskeleton. Tyrosine phosphorylation of the C-terminus of beta-catenin at Tyr142, or the VE-cadherin intracellular domain at Tyr 658/731 by Src family kinases, alters the binding affinity of beta-catenin to VEC (Lagendijk et al., 2015; Poter et al., 2005; Nelson 2004). In addition, VEC was shown to be a critical regulator of TGFß (Rudini et al., 2008) that is also a key regulator of HSPC (Larsson et al., 2003; Yamazaki et al., 2009). Several reports indicate that the canonical Wnt/beta-catenin signaling pathway plays a pivotal role in hematopoietic stem- and progenitor-cell development (Reya et al., 2005; Ruiz Herguido et al., 2012). Interplay between beta-catenin and VE-cadherin may modulate the stem-cell properties of newborn HSPCs. Whether this interaction influences HSPC emergence and embryonic migration and how this interaction contributes to sustained HSPC self-renewal warrant further investigation. Such information may bring us closer to dissecting out the self-renewal mechanisms operating in HSPCs *ex vivo* to further apply these discoveries into clinical applications.

## ACKNOWLEDGEMENTS

We are grateful to Dr André Herbelin for providing key reagents for the experiments and to Sophie Gournet for help in illustrations. We thank the preclinical research platform from the Gustave Roussy Institut for NSG breeding and nursing.

## AUTHOR CONTRIBUTIONS

Contributions: E.O. designed and performed research, analysed data, and wrote the paper; M.T.M.G. performed CFC-assay; D.C. performed cell sorting and contributed to figure preparation; Y.Z. contributed to technical assistance for in vivo experiments and contributed to figure preparation; B.M. and H.B. provided human embryonic and fetal tissues; F.L. contributed to *in vivo* experimental design. A.B.G contributed to discussion of experimental design and results.

## REFERENCES

Arai F, Suda T. Maintenance of quiescent hematopoietic stem cells in the osteoblastic niche. Ann N Y Acad Sci. 2007 Jun;1106:41–53.

Bárcena A, Kapidzic M, Muench MO, Gormley M, Scott MA, Weier JF, Ferlatte C, Fisher SJ. The human placenta is a hematopoietic organ during the embryonic and fetal periods of development. Dev Biol. 2009 Mar 1;327(1):24–33.

Batsivari A, Rybtsov S, Souilhol C, Binagui-Casas A, Hills D, Zhao S, Travers P, Medvinsky A. Understanding Hematopoietic Stem Cell Development through Functional Correlation of Their Proliferative Status with the Intra-aortic Cluster Architecture. Stem Cell Reports. 2017 Jun 6;8(6):1549–1562.

Bertrand JY, Chi NC, Santoso B, Teng S, Stainier DY, Traver D. Haematopoietic stem cells derive directly from aortic endothelium during development. Nature. 2010 Mar 4;464(7285):108–11.

Boisset JC, van Cappellen W, Andrieu-Soler C, Galjart N, Dzierzak E, Robin C. In vivo imaging of haematopoietic cells emerging from the mouse aortic endothelium. Nature. 2010 Mar 4;464(7285):116–20.

Boisset JC, Clapes T, Klaus A, Papazian N, Onderwater J, Mommaas-Kienhuis M, Cupedo T, Robin C. Progressive maturation toward hematopoietic stem cells in the mouse embryo aorta. Blood. 2015 Jan 15;125(3):465–9.

Bowie MB, Kent DG, Copley MR, Eaves CJ. Steel factor responsiveness regulates the high self-renewal phenotype of fetal hematopoietic stem cells. Blood. 2007 Jun 1;109(11):5043–8.

Bowie MB, McKnight KD, Kent DG, McCaffrey L, Hoodless PA, Eaves CJ. Hematopoietic stem cells proliferate until after birth and show a reversible phase-specific engraftment defect. J Clin Invest. 2006 Oct;116(10):2808–16.

Cheifetz S, Bellón T, Calés C, Vera S, Bernabeu C, Massagué J, Letarte M. Endoglin is a component of the transforming growth factor-beta receptor system in human endothelial cells. J Biol Chem. 1992 Sep 25;267(27):19027–30.

Ciau-Uitz A, Patient R. The embryonic origins and genetic programming of emerging haematopoietic stem cells. FEBS Lett. 2016 Nov;590(22):4002–4015.

Chen MJ, Li Y, De Obaldia ME, Yang Q, Yzaguirre AD, Yamada-Inagawa T, Vink CS, Bhandoola A, Dzierzak E, Speck NA. Erythroid/myeloid progenitors and hematopoietic stem cells originate from distinct populations of endothelial cells. Cell Stem Cell. 2011 Dec 2;9(6):541–52.

Chen MJ, Yokomizo T, Zeigler BM, Dzierzak E, Speck NA. Runx1 is required for the endothelial to haematopoietic cell transition but not thereafter. Nature. 2009 Feb 12;457(7231):887–91.

Crisan M, Dzierzak E. The many faces of hematopoietic stem cell heterogeneity. Development. 2016 Dec 15;143(24):4571–4581.

Eilken HM, Nishikawa S, Schroeder T. Continuous single-cell imaging of blood generation from haemogenic endothelium. Nature. 2009 Feb 12;457(7231):896–900.

Ema H, Nakauchi H. Expansion of hematopoietic stem cells in the developing liver of a mouse embryo. Blood. 2000 Apr 1;95(7):2284–8.

Fraser ST, Ogawa M, Yokomizo T, Ito Y, Nishikawa S, Nishikawa SI. Putative intermediate precursor between hematogenic endothelial cells and blood cells in the developing embryo. Dev Growth Differ. 2003 Feb;45(1):63–75.

Fraser ST, Ogawa M, Yu RT, Nishikawa S, Yoder MC, Nishikawa S. Definitive hematopoietic commitment within the embryonic vascular endothelial-cadherin (+) population. Exp Hematol. 2002 Sep;30(9):1070–8.

Gekas C, Dieterlen-Lièvre F, Orkin SH, Mikkola HK. The placenta is a niche for hematopoietic stem cells. Dev Cell. 2005 Mar;8(3):365–75.

Glimm H, Eaves CJ. Direct evidence for multiple self-renewal divisions of human in vivo repopulating hematopoietic cells in short-term culture. Blood. 1999;94:2161–2168.

Itoh K, Tezuka H, Sakoda H, Konno M, Nagata K, Uchiyama T, Uchino H, Mori KJ. Reproducible establishment of hemopoietic supportive stromal cell lines from murine bone marrow. Exp Hematol. 1989;17:145–153.

Ivanovs A, Rybtsov S, Ng ES, Stanley EG, Elefanty AG, Medvinsky A. Human haematopoietic stem cell development: from the embryo to the dish. Development. 2017 Jul 1;144(13):2323–2337.

Ivanovs A, Rybtsov S, Anderson RA, Turner ML, Medvinsky A. Identification of the niche and phenotype of the first human hematopoietic stem cells. Stem Cell Reports. 2014 Mar 27;2(4):449–56.

Ivanovs A, Rybtsov S, Welch L, Anderson RA, Turner ML, Medvinsky A. Highly potent human hematopoietic stem cells first emerge in the intraembryonic aorta-gonad-mesonephros region. J Exp Med. 2011 Nov 21;208(12):2417–27.

Jaffredo T, Gautier R, Eichmann A, Dieterlen-Lièvre F. Intraaortic hemopoietic cells are derived from endothelial cells during ontogeny. Development. 1998 Nov;125(22):4575–83.

Jokubaitis VJ, Sinka L, Driessen R, et al. Angiotensin-converting enzyme (CD143) marks hematopoietic stem cells in human embryonic, fetal, and adult hematopoietic tissues. Blood. 2008;111(8):4055–4063.

Julien E, El Omar R, Tavian M. Origin of the hematopoietic system in the human embryo. FEBS Lett. 2016 Nov;590(22):3987–4001.

Kieusseian A, Brunet de la Grange P, Burlen-Defranoux O, Godin I, Cumano A. Immature hematopoietic stem cells undergo maturation in the fetal liver. Development. 2012 Oct;139(19):3521–30.

Kim I, Yilmaz OH, Morrison SJ. CD144 (VE-cadherin) is transiently expressed by fetal liver hematopoietic stem cells. Blood. 2005 Aug 1;106(3):903–5.

Kissa K, Herbomel P. Blood stem cells emerge from aortic endothelium by a novel type of cell transition. Nature. 2010 Mar 4;464(7285):112–5.

Klaus A, Robin C. Embryonic hematopoiesis under microscopic observation. Dev Biol. 2017 Aug 15;428(2):318–327.

Lagendijk AK, Hogan BM. VE-cadherin in vascular development: a coordinator of cell signaling and tissue morphogenesis. Curr Top Dev Biol. 2015;112:325–52.

Lampugnani MG, Resnati M, Raiteri M, Pigott R, Pisacane A, Houen G, Ruco LP, Dejana E. A novel endothelial-specific membrane protein is a marker of cell-cell contacts. J Cell Biol. 1992 Sep;118(6):1511–22.

Larsson J, Blank U, Helgadottir H, Björnsson JM, Ehinger M, Goumans MJ, Fan X, Levéen P, Karlsson S. TGF-beta signaling-deficient hematopoietic stem cells have normal self-renewal and regenerative ability in vivo despite increased proliferative capacity in vitro. Blood. 2003 Nov 1;102(9):3129–35.

Luckett WP. Origin and differentiation of the yolk sac and extraembryonic mesoderm in presomite human and rhesus monkey embryos. Am J Anat. 1978 May;152(1):59–97.

Majeti R, Park CY, Weissman IL. Identification of a hierarchy of multipotent hematopoietic progenitors in human cord blood. Cell Stem Cell. 2007 Dec 13;1(6):635–45.

Medvinsky A, Rybtsov S, Taoudi S. Embryonic origin of the adult hematopoietic system: advances and questions. Development. 2011 Mar;138(6):1017–31.

Migliaccio G, Migliaccio AR, Petti S, Mavilio F, Russo G, Lazzaro D, Testa U, Marinucci M, Peschle C. Human embryonic hemopoiesis. Kinetics of progenitors and precursors underlying the yolk sac-liver transition. J Clin Invest. 1986 Jul;78(1):51–60.

Morrison SJ, Hemmati HD, Wandycz AM, Weissman IL. The purification and characterization of fetal liver hematopoietic stem cells. Proc Natl Acad Sci U S A. 1995 Oct 24;92(22):10302–6.

Muench MO, Kapidzic M, Gormley M, Gutierrez AG, Ponder KL, Fomin ME, Beyer AI, Stolp H, Qi Z, Fisher SJ, Bárcena A. The human chorion contains definitive hematopoietic stem cells from the fifteenth week of gestation. Development. 2017 Apr 15;144(8):1399–1411.

Nelson WJ, Nusse R. Convergence of Wnt, beta-catenin, and cadherin pathways. Science. 2004 Mar 5;303(5663):1483–7.

Nishikawa SI, Nishikawa S, Kawamoto H, Yoshida H, Kizumoto M, Kataoka H, Katsura Y. In vitro generation of lymphohematopoietic cells from endothelial cells purified from murine embryos. Immunity. 1998(a) Jun;8(6):761–9.

Nishikawa SI, Nishikawa S, Hirashima M, Matsuyoshi N, Kodama H. Progressive lineage analysis by cell sorting and culture identifies FLK1 + VE-cadherin + cells at a diverging point of endothelial and hemopoietic lineages. Development. 1998(b) May;125(9):1747–57.

North TE, de Bruijn MF, Stacy T, Talebian L, Lind E, Robin C, Binder M, Dzierzak E, Speck NA. Runx1 expression marks long-term repopulating hematopoietic stem cells in the midgestation mouse embryo. Immunity. 2002 May;16(5):661–72.

Oberlin E, Fleury M, Clay D, Petit-Cocault L, Candelier JJ, Mennesson B, Jaffredo T, Souyri M. VEcadherin expression allows identification of a new class of hematopoietic stem cells within human embryonic liver. Blood. 2010a Nov 25;116(22):4444–55.

Oberlin E, El Hafny B, Petit-Cocault L, Souyri M. Definitive human and mouse hematopoiesis originates from the embryonic endothelium: a new class of HSCs based on VE-cadherin expression. Int J Dev Biol. 2010b;54(6–7):1165–73.

Oberlin E, Tavian M, Blazsek I, Péault B. Blood-forming potential of vascular endothelium in the human embryo. Development. 2002 Sep;129(17):4147–57.

O’Rahilly R, Muller F, Hutchins GM, Moore GW. Computer ranking of the sequence of appearance of 73 features of the brain and related structures in staged human embryos during the sixth week of development. Am J Anat. 1987;180:69–86.

Ottersbach K, Dzierzak E. The murine placenta contains hematopoietic stem cells within the vascular labyrinth region. Dev Cell. 2005 Mar;8(3):377–87.

Palis J, Yoder MC. Yolk-sac hematopoiesis: the first blood cells of mouse and man. Exp Hematol. 2001 Aug;29(8):927–36.

Palis J, Robertson S, Kennedy M, Wall C, Keller G. Development of erythroid and myeloid progenitors in the yolk sac and embryo proper of the mouse. Development. 1999 Nov;126(22):5073–84.

Potter MD, Barbero S, Cheresh DA. Tyrosine phosphorylation of VE-cadherin prevents binding of p120-and beta-catenin and maintains the cellular mesenchymal state. J Biol Chem. 2005;280:31906–31912.

Prashad SL, Calvanese V, Yao CY, Kaiser J, Wang Y, Sasidharan R, Crooks G, Magnusson M, Mikkola HK. GPI-80 defines self-renewal ability in hematopoietic stem cells during human development. Cell Stem Cell. 2015 Jan 8;16(1):80–7.

Robin C, Bollerot K, Mendes S, Haak E, Crisan M, Cerisoli F, Lauw I, Kaimakis P, Jorna R, Vermeulen M, Kayser M, van der Linden R, Imanirad P, Verstegen M, Nawaz-Yousaf H, Papazian N, Steegers E, Cupedo T, Dzierzak E. Human placenta is a potent hematopoietic niche containing hematopoietic stem and progenitor cells throughout development. Cell Stem Cell. 2009; Oct 2;5(4):385–95.

Reya T, Clevers H. Wnt signalling in stem cells and cancer. Nature. 2005;434:843–850.

Rhodes KE, Gekas C, Wang Y, Lux CT, Francis CS, Chan DN, Conway S, Orkin SH, Yoder MC, Mikkola HK. The emergence of hematopoietic stem cells is initiated in the placental vasculature in the absence of circulation. Cell Stem Cell. 2008 Mar 6;2(3):252–63.

Rudini N, Felici A, Giampietro C, Lampugnani M, Corada M, Swirsding K, Garrè M, Liebner S, Letarte M, ten Dijke P, Dejana E. VE-cadherin is a critical endothelial regulator of TGF-beta signalling. EMBO J. 2008 Apr 9;27(7):993–1004.

Ruiz-Herguido C, Guiu J, D’Altri T, Inglés-Esteve J, Dzierzak E, Espinosa L, Bigas A. Hematopoietic stem cell development requires transient Wnt/β-catenin activity. J Exp Med. 2012 Jul 30;209(8):1457–68.

Rybtsov S, Batsivari A, Bilotkach K, Paruzina D, Senserrich J, Nerushev O, Medvinsky A Tracing the origin of the HSC hierarchy reveals an SCF-dependent, IL-3-independent CD43-embryonic precursor. Stem Cell Reports. 2014 Sep 9;3(3):489–501.

Rybtsov S, Sobiesiak M, Taoudi S, Souilhol C, Senserrich J, Liakhovitskaia A, Ivanovs A, Frampton J, Zhao S, Medvinsky A. Hierarchical organization and early hematopoietic specification of the developing HSC lineage in the AGM region. J Exp Med. 2011 Jun 6;208(6):1305–15.

Shapiro HM. Flow cytometric estimation of DNA and RNA content in intact cells stained with Hoechst 33342 and pyronin Y. Cytometry. 1981;2: 143–150.

Silver L, Palis J. Initiation of murine embryonic erythropoiesis: a spatial analysis. Blood. 1997 Feb 15;89(4):1154–64.

Sinka L, Biasch K, Khazaal I, Péault B, Tavian M. Angiotensin-converting enzyme (CD143) specifies emerging lympho-hematopoietic progenitors in the human embryo. Blood. 2012 Apr 19;119(16):3712–23.

Taoudi S, Gonneau C, Moore K, Sheridan JM, Blackburn CC, Taylor E, Medvinsky A. Extensive hematopoietic stem cell generation in the AGM region via maturation of VE-cadherin + CD45 + predefinitive HSCs. Cell Stem Cell. 2008 Jul 3;3(1):99–108.

Taoudi S, Morrison AM, Inoue H, Gribi R, Ure J, Medvinsky A. Progressive divergence of definitive haematopoietic stem cells from the endothelial compartment does not depend on contact with the foetal liver. Development. 2005 Sep;132(18):4179–91.

Tavian M., Robin C., Coulombel L., and Peault B. The human embryo, but not its yolk sac, generates lympho-myeloid stem cells: mapping multipotent hematopoietic cell fate in intraembryonic mesoderm. Immunity. 2001;15:487–495.

Tober J, Koniski A, McGrath KE, Vemishetti R, Emerson R, de Mesy-Bentley KK, Waugh R, Palis J. The megakaryocyte lineage originates from hemangioblast precursors and is an integral component both of primitive and of definitive hematopoiesis. Blood. 2007 Feb 15;109(4):1433–41.

Wilson A, Laurenti E, Oser G, van der Wath RC, Blanco-Bose W, Jaworski M, Offner S, Dunant CF, Eshkind L, Bockamp E, Lió P, Macdonald HR, Trumpp A. Hematopoietic stem cells reversibly switch from dormancy to self-renewal during homeostasis and repair. Cell. 2008 Dec 12;135(6):1118–29.

Yamazaki S, Iwama A, Takayanagi S, Eto K, Ema H, Nakauchi H. TGF-beta as a candidate bone marrow niche signal to induce hematopoietic stem cell hibernation. Blood. 2009 Feb 5;113(6):1250–6.

Zovein AC, Hofmann JJ, Lynch M, French WJ, Turlo KA, Yang Y, Becker MS, Zanetta L, Dejana E, Gasson JC, Tallquist MD, Iruela-Arispe ML. Fate tracing reveals the endothelial origin of hematopoietic stem cells. Cell Stem Cell. 2008 Dec 4;3(6):625–36.

